# Pyrazinamide kills *Mycobacterium tuberculosis* via pH-driven weak-acid permeation and cytosolic acidification

**DOI:** 10.1101/2025.09.26.678883

**Authors:** Janïs Laudouze, Tatyana I. Rokitskaya, Akira Abolet, Vanessa Point, Alexander M. Firsov, Ljudmila S. Khailova, Maria Sofia Deschutter, Gabriela Gago, Jean-François Cavalier, Stéphane Canaan, Alain R. Baulard, Yuri N. Antonenko, Alexandre Gouzy, Pierre Santucci

## Abstract

Pyrazinamide (PZA) is a cornerstone drug in tuberculosis (TB) treatment with a strong bactericidal activity *in vivo* on both actively and non-replicating bacterial subpopulations. Yet the precise mode of action of its active form, pyrazinoic acid (POA), remains unclear.

In this study, we comprehensively explore and challenge the two major and conflicted models of PZA mode of action. The pH-dependent model, where the drug is mostly effective at acidic pH by acidifying *Mycobacterium tuberculosis* (*Mtb*) cytosol, and the PanD-dependent model where PZA’s active form targets the aspartate decarboxylase PanD, therefore depleting pantothenate (Panto) and subsequently coenzyme A (CoA) levels regardless of the surrounding pH.

By combining standard antimicrobial susceptibility testing at various pH with fluorescence-based live recording of *Mtb* intrabacterial pH (IBpH), we demonstrate that PZA kills *Mtb* by decreasing IBpH, independently of Panto levels. Comparative studies between a prototrophic *Mtb* strain and a Panto auxotrophic mutant lacking the *panCD* locus confirmed that PZA bactericidal activity is primarily driven by pH and its ability to acidify *Mtb* cytosol, independently of the aspartate decarboxylase PanD. Bio-electrophysiology experiments revealed that acidic pH promotes the conversion of the pyrazinoate anion POA⁻ into HPOA which in turn acts as conventional weak acid that facilitates membrane permeation and cytosolic acidification. Finally, using custom culture media, we demonstrate that PZA displays heterogeneous efficacy according to the media composition, therefore proposing a revisited biological model that might explain the discrepancies around PZA’s unique mode of action.

Overall, this work constitutes the first comprehensive side-by-side investigation of the two models and univocally supports a pH-dependent mechanism of action underlying PZA sterilizing activity, providing new insights for the development of more effective PZA-like drugs.

## Introduction

According the WHO’s Global Tuberculosis Report 2025, tuberculosis (TB) remains the leading cause of death worldwide due to an infectious agent with 1.23 million deaths and approximately 10.7 million new cases^1^. For drug-susceptible TB, WHO still recommends the long standing six-month, two-phase regimen that includes a daily cocktail of isoniazid (INH), rifampicin (RIF), ethambutol (EMB), and pyrazinamide (PZA) during 2-months followed by 4 months of RIF and INH. Globally, this regimen achieved a treatment-success rate of 88% for patients who started therapy in 2022, the most recent cohort with complete outcome data ^1^. Despite its effectiveness, taking multiple drugs daily for six months often affects patient compliance which can ultimately lead to treatment failures and the emergence of drug-resistant strains ^1^. Therefore, deepening our understanding of the mechanisms of action of existing anti-TB agents is essential to develop shorter, safer and more potent regimens that improve compliance and curb resistance ^2^.

Among the four front-line anti-TB drugs, PZA holds a singular position. Whereas the molecular target of INH, RIF and EMB are now well-defined, PZA’s mode of action is still actively debated. This is likely due to the fact that PZA is the prototype of a context-dependent drug, that displays very potent efficacy *in vivo* but almost no antibacterial effect in standard mycobacterial broth cultures ^3, 4, 5^. Nevertheless, according to the well-acknowledged Mitchison hypothesis ^6^, its unique *in vivo* sterilizing activity, particularly against non-replicating tubercle bacilli, is what allows the standard of care six-month regimen to prevent relapses ^6, 7^. In that context, PZA remains a key component of most first line current combinations ^8^.

Discovered in 1952, the anti-TB properties of PZA were first reported following extensive synthesis of nicotinamide (NAM) analogues and the subsequent screening of their antibacterial activities within TB-mouse models of infection ^9, 10^. Moreover, the high sterilizing properties of PZA when combined with INH and/or RIF, resulted in its inclusion in the standard anti-TB drug regimen ^11, 12^.

Since its discovery in the 1950’s, seminal studies have demonstrated that PZA is a prodrug that requires to be bio-converted into pyrazinoic acid (POA) to be effective. This activation step is mediated by PncA, a non-essential *Mtb* pyrazinamidase/nicotinamidase involved in the nicotinamide adenine dinucleotide (NAD^+^) salvage pathway ^13, 14^. Consistently, the majority of PZA-resistant clinical isolates harbour mutations in the promoter or the coding region of the gene *pncA* which reduce or abolish PncA production, stability or enzymatic activity, therefore preventing PZA bio-conversion ^15, 16, 17, 18^.

Beyond this obligatory activation step, the downstream molecular events remain contentious. Early proposals, including inhibition of fatty acid synthetase I (FAS I) or the *trans*-translation machinery, did not survive experimental scrutiny ^19, 20, 21, 22^. However, two competing models remain supported by distinct experimental evidence: the pH-dependent weak-acid permeation model and the PanD/CoA-inhibition model.

PZA’s antibacterial activity was originally proposed to be target-independent and driven by the acidic pH that surrounds *Mtb* ^23, 24, 25, 26^. In this first model, PZA diffuses through the mycobacterial cell-wall, and is converted into the negatively charged pyrazinoate anion (POA^-^) by the cytoplasmic PncA enzyme. This negatively charged POA^-^ is further exported extracellularly through an active process that is not yet fully characterised. Then, the protonation state of the HPOA/POA^-^ acid-base pair in the vicinity of the bacterium is dictated by the Henderson-Hasselbalch equation, which relates the pKa of the carboxylic acid function (pKa = 3.6) to the surrounding extrabacterial pH ^24, 26, 27, 28^. Accordingly, highly acidic environment drives the equilibrium towards the neutral protonated HPOA form, which efficiently diffuses through the mycobacterial cell wall and further dissociates into POA^-^ and H^+^ when reaching the circum-neutral bacterial cytosolic pH. The release of protons inside *Mtb* cytoplasm causes a decrease in the intrabacterial pH (IBpH) and membrane potential which has been reported to be critical for PZA antimicrobial activity ^24, 26, 28, 29, 30^.

Nonetheless, this model has been actively challenged over the past decades and an alternative, pH-independent, mode of action has been proposed ^31, 32, 33, 34, 35, 36, 37^. Genomic analysis of *in-vitro* selected *Mtb* low-level resistant mutants to POA revealed mutations in distinct genes including the *panD* gene, suggesting that the aspartate decarboxylase PanD, involved in CoA synthesis, could be the primary target of the drug ^31, 32, 33, 38^. Further biochemical studies have proposed that POA could inhibit PanD activity by binding to its active site ^36^, or alternatively triggering its degradation by the ClpC1-ClpP proteolytic system ^37^. Hence, according to this second model, cytoplasmic POA binds to PanD, limits its activity and induces its degradation, thus ultimately preventing CoA synthesis independently of the extrabacterial pH ^36, 37^.

Given the ongoing debate surrounding the mode of action of PZA, we aimed at comprehensively challenge these two models side-by-side. By combining standard antimicrobial susceptibility testing at various pH with fluorescence-based live recording of *Mtb* IBpH and bio-electrophysiology experiments, we demonstrate that POA alters cytosolic pH of both actively-growing and non-replicating *Mtb* through a weak acid permeation model. Our study reports that this process was only minimally affected by Pantothenate (Panto) and β-alanine (β-ala) levels. Moreover, genetic studies, using a *panCD* mutant strain, confirmed that PZA bactericidal action is primarily driven by acidic pH and its ability to acidify the *Mtb* cytosol, independently of the aspartate decarboxylase PanD. Finally, by using custom culture media containing a physiologically more relevant lipid-based carbon source, we demonstrate that the antagonistic effect of the CoA precursors observed in standard 7H9 media are no longer detectable. These results not only shed new light on the mode of action of PZA but also explain some of the long-standing discrepancies between its observed *in vitro* and *in vivo* activities.

## Results

### Comprehensive pH-dependent analysis of PZA/POA antibacterial activity

Since the pH-dependent mode of action of PZA/POA has recently been challenged in our community ^39, 40, 41, 42^, we first aimed to re-evaluate the impact of mildly acidic pH on the efficacy of a panel of anti-TB drugs including PZA/POA (Fig 1a). Antimicrobial susceptibility testing was then performed in standard Middlebrook 7H9 medium at pH 6.8 and pH 5.5, and the minimal inhibitory concentration (MIC) against *Mtb* H37Ra was determined for each compound (Fig 1b). As expected, PZA was inactive when assessed at pH 6.8 with MIC values exceeding 500 μg/mL, while POA displayed a very moderate activity with an MIC of 125 μg/mL. As previously reported ^24, 25^, both PZA and POA exhibited significant pH-dependent growth inhibition patterns with an approximate 4 to 8-fold decrease in MIC values when tested at pH 5.5 (Fig 1b). This pH-dependent effect was also easily visualised in a heatmap using color-coded representation based on Log^2^(FC_MIC_) values (Fig 1d). Additional experiments with the natural substrate and product of PncA, nicotinamide (NAM) and nicotinic acid (NAC) respectively, showed similar inhibition patterns with at least a 4-fold gain in potency when tested at acidic pH, thereby reflecting a pH-dependent activity (Fig 1b). As previously observed, salicylic acid (SA), used in control experiment, showed a strong potency at acidic pH with a significant shift from 200 μg/mL to 25 μg/mL ^43, 44^. By contrast, no pH-dependent shift in MIC was observed with isoniazid (INH), ethionamide (ETH) and prothionamide (PTH), three well-established pro-drugs that target the 2-trans-enoyl-ACP reductase InhA (Fig 1b). This confirms that the observed pH-dependent activity is compound and/or mode of action dependent rather than universal. Altogether, these results support a model in which an acidic pH environment potentiates the activity of several anti-TB drugs or prodrugs bearing carboxylic acid function, conferring weak acid properties, including PZA and POA.

**Figure 1.**
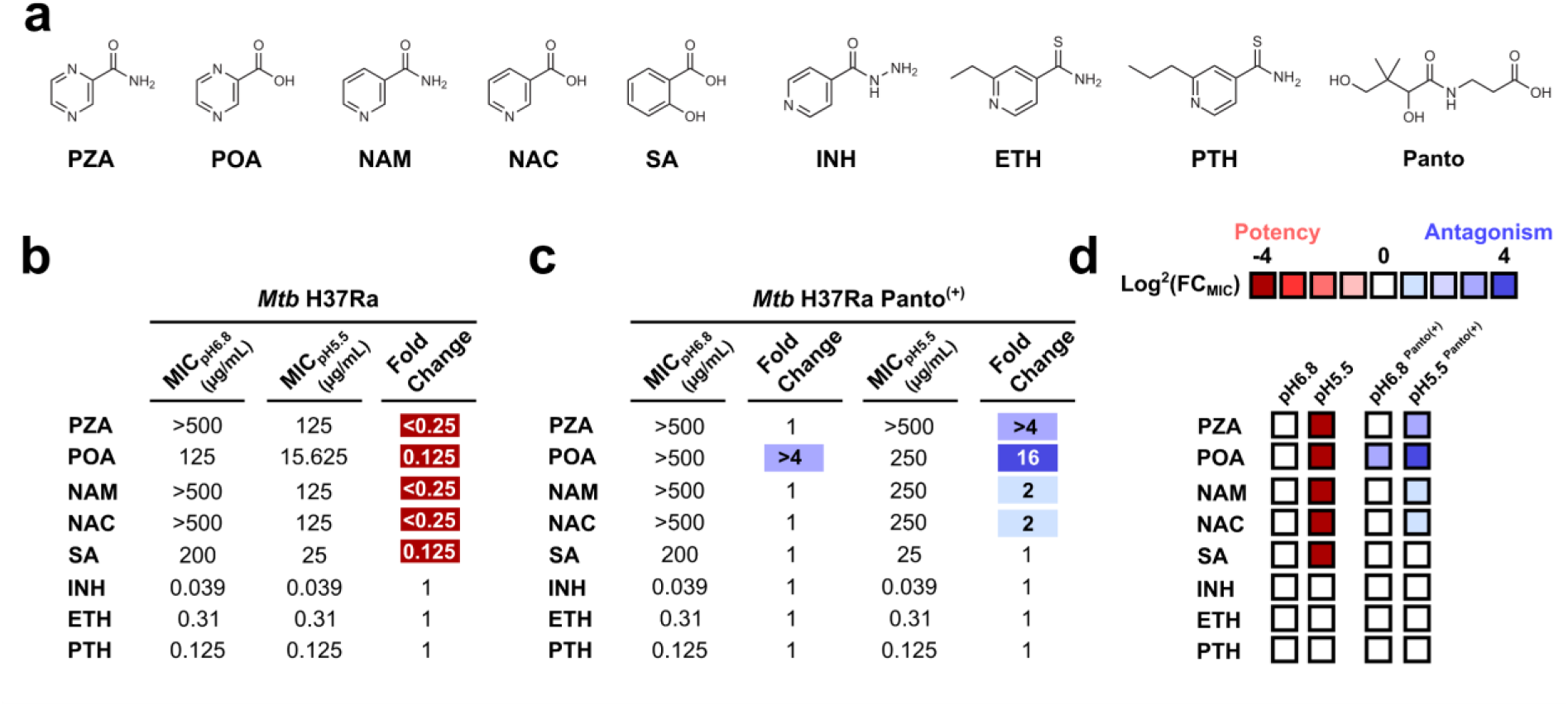
Comprehensive analysis of PZA/POA antibacterial activity against *Mtb*. **(a)** Structure of the compounds investigated in this study. The following compounds were used Pyrazinamide (PZA); Pyrazinoic acid (POA); Nicotinamide (NAM); Nicotinic acid (NAC); Salicylic acid (SA); Isoniazid (INH); Ethionamide (ETH); Prothionamide (PTH) and Pantothenate (Panto) **(b)** *Mtb* antibiotic susceptibility profiling performed at circum-neutral (pH6.8) and acidic (pH5.5) pH in the absence of Panto. Results are expressed as minimal inhibitory concentration (MIC). The fold change in MIC values is expressed as by applying the following formula MIC_pH5.5_/MIC_pH6.8_. **(c)** *Mtb* antibiotic susceptibility profiling performed at circum-neutral (pH6.8) and acidic (pH5.5) pH in the presence of 25 µg/mL of Panto (Panto^(+)^). Results are expressed as MIC. The fold change in MIC values is expressed as by applying the following formula MIC_Antagonist_/MIC_CTRL_ for a given pH of pH6.8 or pH5.5. **(d)** Potency and antagonisms determined from **(b)** or **(c)** are further displayed as color-coded heat map from red (potency) to blue (antagonism). Heat map intensities for each compound represent the fold change (FC) in MIC values expressed as Log^2^(FC_MIC_) by applying the following formula MIC_pH5.5_/MIC_pH6.8_ or MIC_Antagonist_/MIC_CTRL_ for a given pH of pH6.8 or pH5.5. All results are representative of three independent technical replicates performed in three biological replicates.

### Supplementation with CoA precursors downstream of the aspartate decarboxylase PanD antagonizes PZA/POA activity in standard Middlebrook 7H9 medium

The aspartate decarboxylase PanD catalyzes the decarboxylation of L-aspartate (L-Asp) to form β-alanine (β-ala), involving the removal of a carboxyl group, the consumption of a proton (H⁺), and the release of CO₂. Subsequently, the pantothenate synthetase (PanC) catalyzes the ATP-dependent condensation of β-Ala with pantoate, yielding Panto, a vital precursor for the biosynthesis of CoA via the CoaABCDE proteins (Fig S1).

One of the main pieces of evidence supporting PanD as the primary target of PZA/POA was based on the observation that supplementation of complete Middlebrook 7H9 medium with CoA-biosynthesis precursors, such as Panto or β-ala, negatively impacted the efficacy of PZA/POA at pH 6.8 and pH 5.5 ^32, 45^. To confirm these results, we performed a comprehensive analysis of *Mtb*-growth inhibition profiles with our subset of drugs at both circumneutral or mildly acidic pH in the presence of exogenous Panto. In good agreement with literature data, the presence of 25 μg/mL of Panto strongly antagonised PZA/POA efficacy at both pH, resulting in an increase in their MIC values from >4-to 16-fold (Fig 1c & Fig 1d). Interestingly, this modulator only displayed a very weak antagonistic effect at acidic pH against NAM/NAC (Fig 1c & Fig 1d), in line with previous observations ^45^. Finally, no impact on the anti-TB drugs INH, ETH and PTH was observed, nor with the weak acid control molecule SA (Fig 1c & Fig 1d). All these results confirm the multiple observations from independent groups, supporting the concept that supplementation with Panto specifically antagonise PZA/POA efficacy in complete Middlebrook 7H9 media.

### *panCD* locus and CoA-precursors supplementation have minimal effect on PZA/POA-induced intrabacterial pH disruption on replicating *Mtb*

We and others have previously proposed that PZA/POA efficacy was directly linked to its ability to alter *Mtb* IBpH homeostasis in a pH-dependent manner ^24, 29, 30^. To confirm these results, we capitalized from a well-established *Mtb* reporter strain that produces a ratiometric pH-GFP sensor, enabling the non-invasive monitoring of IBpH changes during drug treatment (Fig S2) ^29, 30, 43, 46, 47^. Using this genetic tool, we set up an *in vitro* assay and calibrated the response of this fluorophore using lysates from recombinant *Mtb* H37Ra pH-GFP reporter strain at different pH increments (Fig S2a; top panel). Locally estimated scatterplot smoothing (LOESS) curve fitting was performed in order to obtain a reliable standard curve that links *Mtb* pH-GFP fluorescence ratio to IBpH values (Fig S2a; bottom panel). To validate that this experimental system fully enables the quantitative monitoring of compound-mediated IBpH perturbation *in vitro*, the protonophore carbonyl cyanide *m*-chlorophenyl hydrazone (CCCP) and the pH-dependent ionophore monensin (MON) were used as positive controls (Fig S2b). Exponentially growing *Mtb* pH-GFP was pulsed for 24 h in complete Middlebrook 7H9 medium with increasing concentrations of CCCP (Fig S2b; top panel) or MON (Fig S2a; bottom panel) at circumneutral and acidic pH, respectively. Results showed that both compounds sharply decreased *Mtb* pH-GFP ratio in a dose-dependent manner until reaching IBpH values lower than IBpH 5.5, therefore validating our experimental approach. Control experiments were also performed at pH 6.8 or pH 5.5 in complete 7H9 media in presence/absence of high exogenous Panto levels (*e.g.,* 25 μg/mL) and showed no major changes (*e.g.,* pH-GFP ratios values in between 0.8 and 0.9 in all conditions tested), approximating a constant IBpH value of IBpH 6.5, in line with previous reports ^29, 30, 43, 46, 47^ (Fig S3a).

Dose response analysis upon PZA treatment confirmed previous observations that reported no major cytosolic acidification when assessed at circumneutral pH (Fig 2a; left panel). Conversely, at pH 5.5 PZA strongly impairs *Mtb* IBpH homeostasis, supporting a pH-dependent effect (Fig 2a; central panel). Importantly, these changes were deemed of statistical significance from 31.3 μg/mL, a concentration that is commonly observed in plasma and lungs of TB patients undergoing chemotherapy ^48, 49, 50^.

**Figure 2.**
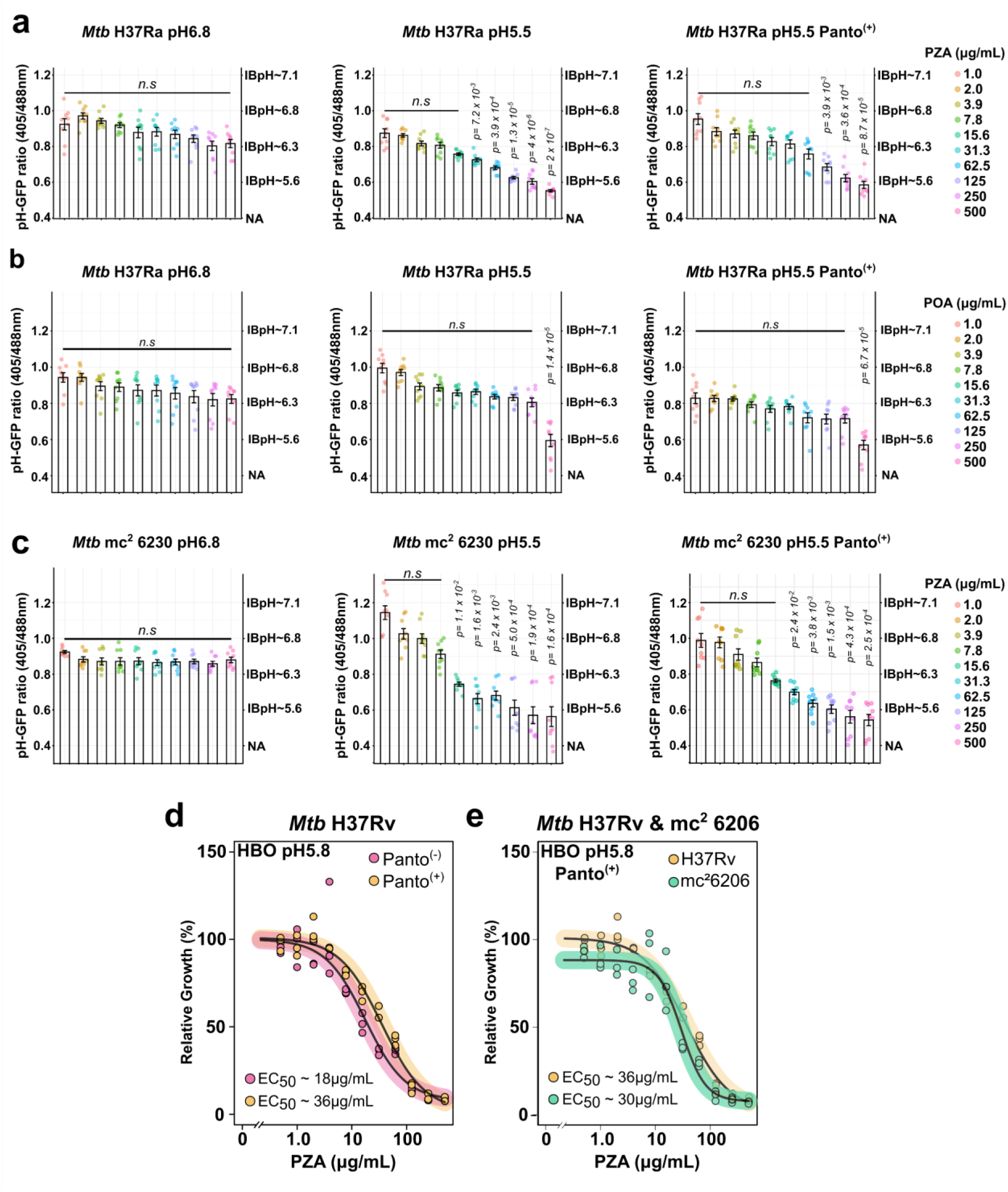
PZA/POA displays pH-dependent antibacterial activity on actively growing *Mtb* by primarily impacting IBpH homeostasis. **(a)** Quantification of PZA-induced *Mtb* H37Ra IBpH alteration at circum-neutral (*left* panel), acidic pH (*middle* panel) and acidic pH in the presence of 25 μg/mL of Panto (*right* panel). *Mtb* pH-GFP ratio were determined in the presence of increasing concentrations of PZA after 24 h of exposure. **(b)** Quantification of POA-induced *Mtb* H37Ra IBpH alteration at circum-neutral (*left* panel), acidic pH (*middle* panel) and acidic pH in the presence of 25 μg/mL of Panto (*right* panel). *Mtb* pH-GFP ratio were determined in the presence of increasing concentrations of POA after 24 h of exposure. **(c)** Quantification of PZA-induced *Mtb* mc^2^ 6230 IBpH alteration at circum-neutral (*left* panel), acidic pH (*middle* panel) and acidic pH in the presence of 25 μg/mL of Panto (*right* panel). *Mtb* pH-GFP ratio were determined in the presence of increasing concentrations of PZA after 24 h of exposure. *Mtb* IBpH results displayed in this figure were obtained from 3 biologically independent experiments and are displayed as mean ± SEM. In dose response analysis, statistical significance was assessed by comparing the means of each concentration with the lowest concentration tested using one-way ANOVA followed with Tukey’s multiple comparisons test. All *p*-values were considered significant when *p*-value < 0.05. **(d)** Comparative analysis of PZA-mediated growth inhibition against *Mtb* H37Rv in HBO medium pH 5.8 and in the absence (pink) or presence of Panto (yellow) antagonist. Dose-response analysis displayed were obtained following a 4-parameter nonlinear logistic regression and EC_50_ were determined accordingly. Results are representative of three biological replicates. **(e)** Comparative analysis of PZA-mediated growth inhibition against *Mtb* H37Rv WT (yellow) and *Mtb* H37Rv mc^2^ 6206 (green) in HBO medium pH 5.8 in the presence of 25 µg/mL Panto antagonist. Dose-response analysis displayed were obtained following a 4-parameter nonlinear logistic regression and EC_50_ were determined accordingly. Results are representative of three biological replicates.

Similar results were obtained regarding *Mtb* IBpH when treated with the active form POA but only at high concentration (Fig 2b; left & central panels), while control experiments with INH did not show any effect on *Mtb* IBpH, whatever the pH and the tested concentrations (Fig S3b). According to the literature, we speculated that the lack of effect of POA could be explained by its poor solubility and increased binding affinity to bovine serum albumin (BSA) contained in the medium in comparison to PZA ^25^. To test the latter hypothesis, we performed antimicrobial susceptibility testing and IBpH monitoring in custom Middlebrook 7H9 broth in the presence or absence of 5 g/L of BSA (Fig S4). Results showed that the presence of BSA had no effect on POA efficacy (Fig S4a) nor its ability to acidify *Mtb* H37Ra cytosolic pH (Fig S4c). To further support our model and demonstrate that PZA-induced IBpH acidification is mediated by POA, we monitored IBpH levels of *Mtb* H37Rv WT, H37Rv Δ*pncA* and Δ*pncA::pncA* recombinant strains (Fig S5). Quantitative analysis of the three strains demonstrated that IBpH acidification requires a functional PncA, since the IBpH of the Δ*pncA* mutant strain is not impacted by PZA treatment. This phenotype was fully reverted while complementing the mutant strain with a functional copy of the *pncA* gene (Fig S5a). Altogether, these results confirm that PncA is essential for PZA-mediated IBpH acidification by converting PZA into its active form POA, further supporting a model in which POA is the major driver of IBpH acidification under acidic conditions.

Altogether, these results, which agree with previous reports ^29, 30^, show that *Mtb* growth arrest mediated by antibiotics at mildly acidic pH is not necessarily the result of IBpH disruption ^43^; and that IBpH perturbation is observed only with a specific subset of drugs, including PZA/POA.

In order to establish whether the recorded IBpH homeostasis perturbations were linked to the PanD-dependent mode of action, and therefore CoA-biosynthesis defect, the same subset of experiments was repeated in the presence of 25 μg/mL Panto (Fig 2a; right panel & Fig S5b), a high exogenous concentration known to fully rescue the growth of an auxotrophic *Mtb panCD* mutant (i.e., *Mtb* mc^2^ 6230 strain) ^51, 52^. In this context, Panto supplementation had a mild but detectable and statistically significant effect on PZA -mediated IBpH perturbation (Fig 2a; right panel). Indeed, in the presence of Panto, 125 μg/mL of PZA where required to trigger significant IBpH acidification, as compared to 31.3 μg/mL in its absence. This result is in line with the negative effect of Panto on PZA/POA efficacy observed during antimicrobial susceptibility profiling assays.

Following up on this observation, antimicrobial susceptibility profiling experiments were performed in the presence of exogenous L-Asp and β-ala, the substrates and products of PanD, respectively. Results displayed in Fig S6, confirmed that β-ala displays negative effects onto PZA/POA efficacy, similar to the ones observed when supplementing with Panto, while no major effects were observed when supplementing with L-Asp. In addition, live monitoring of *Mtb* IBpH in the presence of exogenous β-ala, did not rescue PZA-mediated IBpH acidification, in agreement with the very modest effect of Panto supplementation observed in Fig 2a.

To go further, we decided to complement this series of experiments by relying on a genetic approach. Hence, we tested whether *panCD* deletion could interfere with PZA/POA-mediated IBpH homeostasis perturbation. We first confirmed that the *Mtb* mc^2^ 6230 strain was auxotroph for Panto in our experimental system (Fig S7a). Then, we introduced the pUV15-pHGFP vector in this genetic background to generate a *Mtb* mc^2^ 6230 pH-GFP reporter strain. As for its prototrophic counterpart, IBpH remained constant in *Mtb* mc^2^ 6230 at both pH 6.8 and pH 5.5 and in the presence or absence of Panto (Fig S7b). The observation that *Mtb* mc^2^ 6230 is able to maintain its IBpH in the absence of Panto (bacteriostatic condition) implies that CoA auxotrophy does not trigger major IBpH maintenance defects at least within the first 24h.

Dose-response analysis of PZA/POA impact on IBpH homeostasis using the same experimental setting as above showed no major changes in *Mtb* mc^2^ 6230 IBpH when assessing PZA or its active counterpart POA at circumneutral pH (Fig 2c; left panel & Fig S7c left panel). Interestingly, at pH 5.5, a statistically significant increase in cytosolic acidification caused by PZA was observed starting from 15.6 μg/mL (Fig 2c; central panel). Finally, experiments performed in the presence of 25 μg/mL of Panto showed a very minor antagonistic effect on PZA/POA-mediated acidification (Fig 2c; right panel & Fig S7c).

Because *Mtb* H37Ra and mc²6230 originate from different genetic backgrounds, we performed an additional comparative experiment using a set of strains derived from the same parental background. Specifically, we evaluated H37Rv WT, H37Rv Δ*panCD*, and the complemented H37Rv Δ*panCD*::*panCD* strains, all expressing the pH-GFP reporter. (Fig S8). As showed in Fig S8, the three strains displayed similar PZA-mediated IBpH disruption profiles (Fig S8a), in the presence or absence of exogenous Panto (Fig S8b), confirming our results obtained previously with *Mtb* H37Ra and mc^2^ 6230.

To better characterise the role of Panto and PanD in PZA/POA mode of action, we tested whether the enzymatic reactions carried out by PanC and PanD and/or their down-products might help to mitigate PZA-triggered IBpH acidification, but only at relatively low concentrations of PZA. As such, *Mtb* H37Ra pH-GFP strain was transformed by the L5-integrative pMV306 empty vector ^53^ or its counterpart that contains the *panCD* locus under the control of the strong constitutive promoter Psmyc ^54^ (Fig S9a). Recombinant clones where selected and immune-detection of mature PanD was achieved using anti-His probe (Fig S9b). Quantitative analysis of PZA-mediated IBpH disruption in both pMV306 or pMV306-Psmyc-*panCD* did not reveal any major change (Fig S8c & S8d). Further analysis performed in the presence or absence of PanD substrate or product (i.e. L-Asp or β-ala, respectively), showed that *panCD* overexpression does not prevent PZA-mediated IBpH acidification even by supplementing with precursors from the CoA biosynthesis pathways (Fig S8c & S8d).

Altogether, our data reveal that, even though the deletion of *panCD* and Panto supplementation alter PZA-mediated IBpH disruption to some extent, their impact is minor. Indeed, the disruption of IBpH homeostasis does not appear to depend directly on PanD function or its downstream CoA biosynthetic pathway. Instead, these IBpH alterations occur primarily in a pH-dependent manner and are independent of CoA deficiency or availability of its precursors. Therefore, our data strongly indicate that PanD is not the primary target of PZA/POA.

### Media composition modulates CoA precursors antagonism on PZA/POA efficacy against *Mtb*

It is well acknowledged that media composition, including carbon-sources, can significantly impact the efficacy of antibiotics *in vitro* ^55, 56^. With regards to PZA/POA, Gopal *et al*. showed that an increasing concentration of glycerol *in vitro* hypersensitises *Mtb* to POA further highlighting that the nature but also the availability of carbon source might alter drug efficacy^33^. To confirm this observation, we first compared *Mtb* H37Ra susceptibility profile towards POA at mildly acidic in standard 7H9 media containing 10% OADC in presence or absence of 0.2% glycerol (Fig S10). The obtained results confirm that the presence of this non-physiological concentration of glycerol ^57^ enhances POA efficacy, with EC_50_ values shifting from 38 μg/mL to 9 μg/mL and MIC values from 500 μg/mL to 15.6 μg/mL in glycerol-containing media (Fig S10). Similar significant changes in MIC values, shifting from >500 μg/mL to 125 μg/mL, were also observed with PZA in the presence of glycerol; whereas no alteration in MIC values was reached with the control drug SA (Fig S10c). Remarkably, the presence of glycerol significantly increases PZA/POA efficacy (Fig S10). From these findings, we further investigated how carbon sources alter PZA/POA efficacy. Since the Middlebrook 7H9 media supplemented with OADC contains both glycolytic and lipid carbon sources, we opted for custom minimal media of perfectly controlled and known composition. We capitalized from the recently developed High-BSA-Oleate (HBO) medium, that rely on high-BSA content and oleic acid a sole carbon source to assess PZA activity ^58^. Using this experimental system at pH 5.8, we confirmed that *Mtb* H37Rv reference strain was not only susceptible to PZA (EC_50_ ∼18 μg/mL), but also that the negative effect of Panto was almost completely lost, with EC_50_ ∼36 μg/mL, respectively (Fig 2d). Since Panto had a very limited effect on PZA efficacy in this custom media, we took advantage of this phenotype and perform susceptibility profiling of a *Mtb* H37Rv WT and its respective Δ*panCD* mutant. (Fig 2e). Comparative analysis of the susceptibility profile of WT and Δ*panCD* mutant in presence of Panto showed no major differences, with EC_50_ values of 36 μg/mL and 30 μg/mL, respectively. In this more physiologically relevant medium, our results clearly demonstrate that PZA efficacy is independent of Panto supplementation or the presence of a functional *panCD* locus.

### PZA/POA-mediated pH-disruptive effect against non-replicating *Mtb* is driven by the availability of surrounding protons

According to the Mitchison hypothesis, the sterilizing activity of PZA is greatly enhanced against non-replicating bacteria ^27, 59^. This was confirmed by independent groups that reported potent bactericidal activity *in vitro* when assessed on starved non-replicating *Mtb* ^25, 35, 60^. To go further in the characterisation of PZA’s mode of action, we investigated PZA-mediated IBpH disruption within the well-established phosphate-citrate buffer. This minimalist experimental system enables to test compound-mediated IBpH homeostasis perturbation, at different pH increments on non-replicating starved bacteria ^29, 46^. Hence, we first monitored *Mtb* H37Ra adaptation to phosphate citrate buffer after 48 h of incubation at different pH ranging from pH 5.5 to pH 7 (Fig S11). Monitoring of IBpH revealed that, as pH decreases, *Mtb* H37Ra slightly equilibrates its IBpH towards more alkaline values. This phenomenon is likely a compensatory mechanism in response to protons influx and changes in membrane potential^61^. Interestingly, such pH-dependent adaptation in IBpH was not detectable in standard supplemented Middlebrook 7H9 medium (Fig S3a), suggesting that IBpH homeostasis might be modulated by multiple exogenous factors including nutrients availability during pH variations.

Such approach was further used to monitor the effect of several drugs on *Mtb* H37Ra IBpH homeostasis at various pH, with RIF, and MON as negative and positive controls respectively (Fig S12a-b). First, dose response analysis of PZA treatment showed a clear dose-dependent and pH-dependent effect on *Mtb* H37Ra IBpH (Fig 3a). Similar results were also obtained with NAM/NAC (Fig S12c-d) in agreement with their previously observed pH-dependent potency. Additional control experiments with PTH, ETH and INH (Fig S12e-g) confirmed once again that such pH-dependent activity was specific to PZA/POA and its analogues. When carried out with the *Mtb* mc^2^ 6230 mutant strain, results showed that the lack of aspartate decarboxylase PanD had only a minimal impact on the pH-dependent IBpH-disruptive effect PZA (Fig S13a) or POA (Fig S13b & S13c) in comparison to its prototrophic *Mtb* H37Ra counterpart.

**Figure 3.**
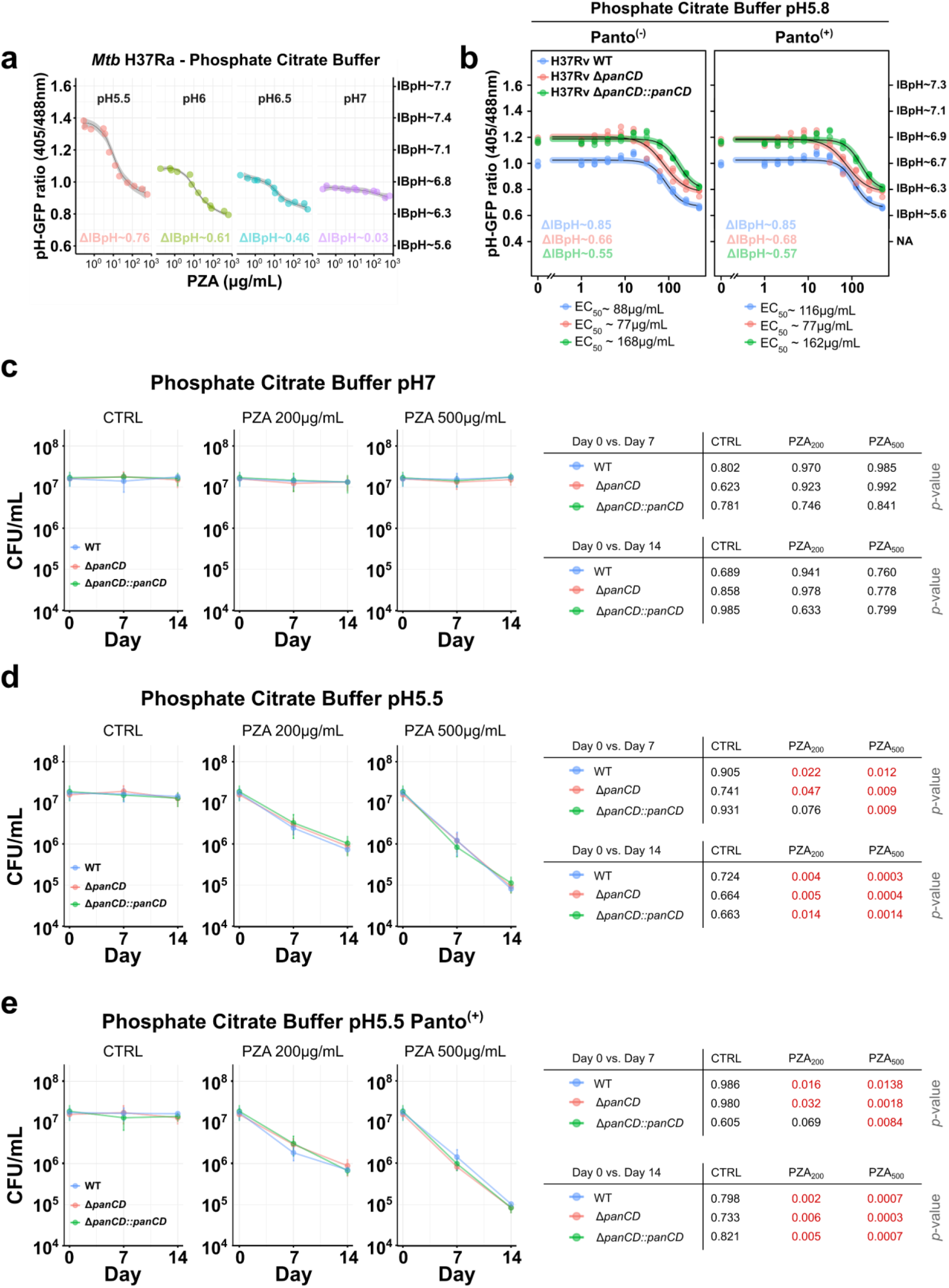
Proton availability dictates the ability of PZA/POA to acidify the cytosol and kill non-replicating *Mtb* independently of the *panCD* locus and pantothenate levels. **(a)** pH-dependent PZA-mediated IBpH cytosolic acidification of *Mtb* H37Ra. Dose-response analysis was performed in phosphate citrate buffer adjusted at pH 7 (purple), pH 6.5 (blue), pH 6 (green) and pH 5.5 (red). *Mtb* were exposed to increasing concentrations of PZA from 0.25 to 500 μg/mL for 48 h before pH-GFP ratio recording. Using pH-GFP ratios, distinct models were built using the LOESS function and ΔIBpH were calculated by comparing the IBpH recorded at the highest PZA concentration tested with the one at the lowest PZA concentration included in the dose response for each tested pH. **(b)** Quantification of PZA-induced IBpH alteration at acidic pH in *Mtb* H37Rv WT (blue), *Mtb* H37Rv Δ*panCD* (pink) and *Mtb* H37Rv Δ*panCD::panCD* (green). Dose-response analyses were performed in phosphate citrate buffer adjusted at pH 5.8, and further obtained following a 4-parameter nonlinear logistic regression and EC_50_ were determined accordingly. *Mtb* were exposed to increasing concentrations of PZA from 0.25 to 500 μg/mL for 48 h before pH-GFP ratio recording. Using pH-GFP ratios, distinct models were built using the LOESS function and ΔIBpH were calculated by comparing the IBpH recorded at the highest PZA concentration tested with the one at the lowest PZA concentration included in the dose response for each tested pH. **(c-d-e)** pH-dependent bactericidal activity of PZA on non-replicating *Mtb* H37Rv WT (blue), *Mtb* H37Rv Δ*panCD* (pink) and *Mtb* H37Rv Δ*panCD::panCD* (green). Survival in phosphate citrate buffer was performed by pulsing 1 x 10^7^ *Mtb* CFU/mL with increasing concentration of PZA including 0 µg/mL, 200 µg/mL and 500 µg/mL in the presence or absence of 25 µg/mL Panto. At day 0, 7 and 14, *Mtb* cells were recovered, serially diluted and plated onto 7H11 agar media in the presence 25 µg/mL of Panto. Plates were further incubated 14-21 days at 37°C before performing CFU determination. Results are representative of three biological replicates. Statistical was assessed after checking variance homogeneity in between two groups of interest and a Student Two Sample t-test was applied. Significant *p*-values are highlighted in red on the right of each panel.

To confirm these results, we then repeated these experiments using *Mtb* H37Rv WT, H37Rv Δ*panCD* and Δ*panCD::panCD* recombinant strains expressing the pH-GFP reporter at acidic pH. As displayed in Fig 3b, some very modest changes were observed in between the three strains with EC_50_ estimated around 88 µg/mL, 77 µg/mL and 168 µg/mL respectively. Supplementation with exogenous Panto did not result in important changes with estimated EC_50_ of 116 µg/mL, 77 µg/mL and 162 µg/mL. Finally, analysis of ΔIBpH revealed that PZA-mediated effect is independent of the PanCD locus and or the level of Panto.

Taken together, these data confirm that PZA/POA can disrupt IBpH in non-replicating *Mtb* in a strictly pH-dependent manner, independent of PanD or CoA biosynthesis. These findings reinforce the idea that the availability of external protons is the critical driver of PZA/POA-mediated pH disruption in non-replicating bacilli.

### Bactericidal activity of PZA on non-replicating *Mtb* is independent of *panCD* locus or pantothenate level

Since both prototrophic and auxotrophic strains display similar non-replicating behaviour in phosphate citrate buffer, and that PZA treatment at acidic pH resulted in important cytosolic acidification for all the strains tested, we investigated whether such acidification could result in bacterial cell death independently of the *panCD* locus.

To do so, exponentially growing *Mtb* H37Rv WT, *Mtb* H37Rv Δ*panCD* and *Mtb* Δ*panCD::panCD* strains were incubated in phosphate citrate buffer at pH 7 or pH 5.5 in the absence or the presence of increasing concentrations of PZA. Samples were collected at day 0, 7 and 14, serially diluted and plated on complete agar media to perform quantitative CFU-based assessment of bacterial viability (Fig 3c-e). A first experiment performed at pH 7 confirmed that PZA treatment had no effect on *Mtb* viability over the 14-days period even at high PZA concentration of 500 µg/mL, confirming once again that PZA-mediated lethality is pH-dependent (Fig 3c). Importantly, the absence of PanC and PanD activity (in the Δ*panCD* mutant) had no effect on *Mtb* viability over this time period, even in the absence of Panto supplementation, supporting the idea that Panto auxotrophy is bacteriostatic rather than cidal in agreement with our data and previous report ^62^.

Subsequently, this experiment was performed at pH 5.5 (Fig 3d). A drug-free control condition was included as reference baseline and demonstrated that *Mtb* viability was not altered during the entire course of the experiment at pH 5.5 regardless the presence or absence of the *panCD* locus (Fig 3d; left panel). Supplementation of 25 μg/mL of Panto had also no impact on *Mtb* viability over the 14-day period (Fig 3e; left panel). On the other hand, PZA treatment showed, as expected, a time and dose dependent effect on bacterial viability. Indeed, treatment with 200 or 500 µg/mL PZA induced approximately 2 to 3-log^10^ reduction in bacterial viability at day 14 for all the tested strains. This reduction in comparison to day 0 was statistically significant for all the strains tested. Nevertheless, no significant differences were observed between *Mtb* H37Rv WT, *Mtb* H37Rv Δ*panCD* and *Mtb* Δ*panCD::panCD* strains. Finally, supplementation of the phosphate citrate buffer with 25 μg/mL Panto had no effect on PZA-mediated cell death, suggesting that PZA sterilizing activity is completely independent of Panto level.

All these results were confirmed using spot-based dilution assays with the alternative strains *Mtb* H37Ra and mc^2^ 6230, where the *panCD* auxotrophic mutant strain exhibited a vulnerability pattern similar to the *Mtb* H37Ra prototrophic strain, showing noticeable time- and dose-dependent effects which resulted in strong viability loss, independently of the presence of Panto (Fig S14).

Altogether, these data univocally support a pH-dependent bactericidal mode of action of PZA that does not require the targeting of PanD nor the disruption of the CoA biosynthesis pathway.

### Bio-electrophysiology experiments reveal that POA pH-disruptive activity is mediated by a weak acid permeation model

PZA/POA-mediated alterations of IBpH homeostasis may arise from two primary mechanisms. First, HPOA, the protonated form of POA, acts as a weak acid, that passively diffuses across the cell envelope from a lower pH compartment to a higher pH compartment. Thus, it gets deprotonated and releases a proton (H+) in a pH-dependent manner, based on its pKa and the cytosolic pH according to well-established Henderson-Hasselbalch equation. Alternatively, POA might exhibit a protonophore-like activity, shuttling protons across the cytoplasmic membrane through repeated transmembrane cycles.

In this context, we first examined the protonophoric activity of PZA and its active form POA in a pure bilayer lipid membrane (BLM) system and further measured the electric current across a planar lipid bilayer under voltage-clamp conditions (Fig 4a). As shown in Fig 4b-c, neither PZA nor POA induced a transmembrane electric current on BLM when assessed at pH 7. Subsequent addition of the well-established protonophore CCCP led to the induction of the BLM current, as expected for a protonophore in this experimental system. Therefore, our results suggest that PZA and POA are not protonophore because they are unable to induce an electrical current in a pure system of planar bilayers.

**Figure 4.**
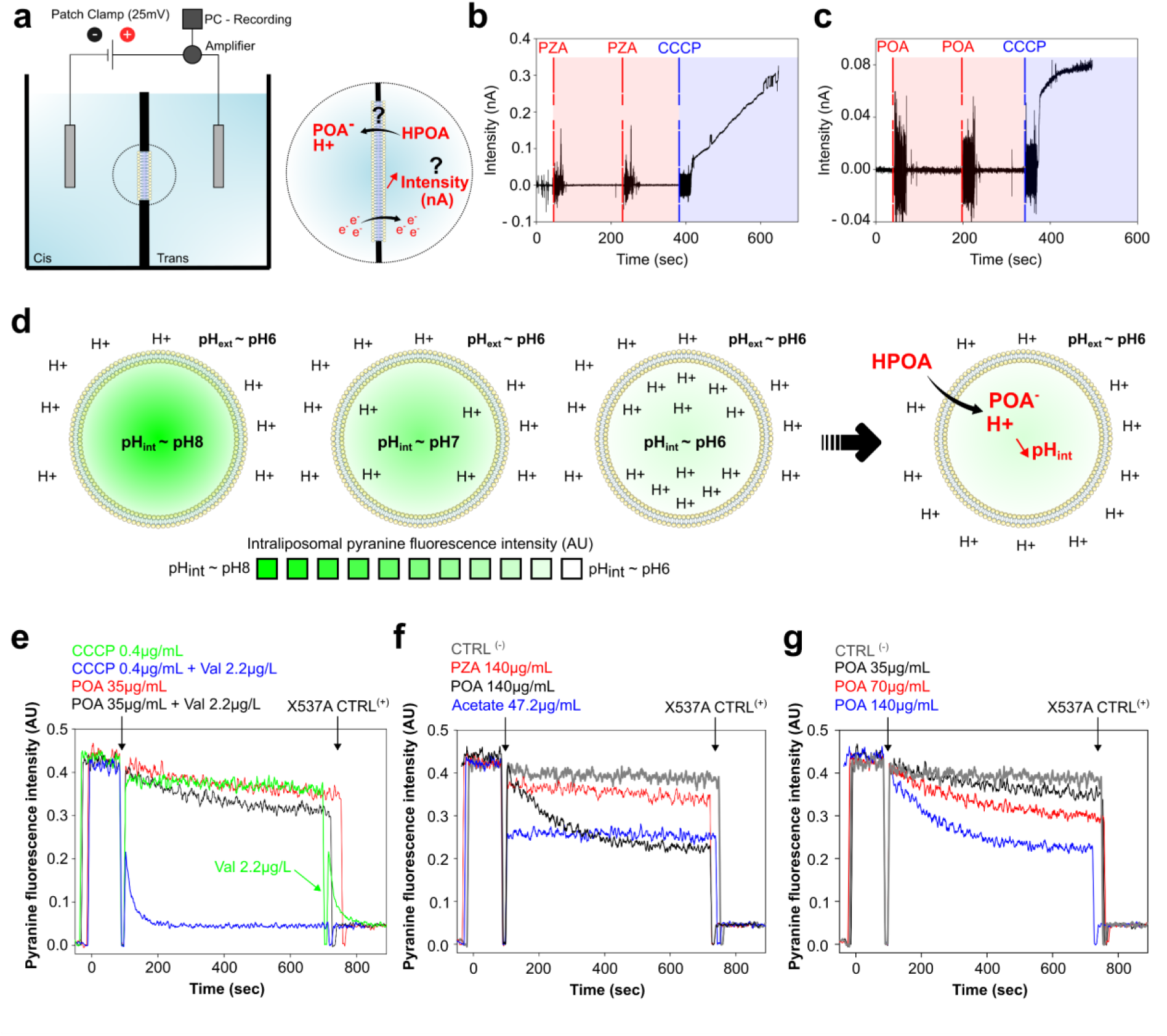
Bio-electrophysiology experiments support a weak acid model of membrane permeation and acidification by the active form POA. **(a)** Schematic representation of the BLM (made from DPhPC) electrophysiological setup used to test the protonophoric activity of PZA and POA. **(b-c)** PZA (panel **b**) or POA (panel **c**) does not induce electrical current through planar BLM made from DPhPC. The well-established protonophore CCCP was used as positive control. Experiments were carried out in a solution composed of 10 mM Tris, 10 mM MES, 100 mM KCl, pH 7.0. BLM voltage was set constant at 50 mV. **(d)** Schematic representation of the pyranine-loaded liposomes setup used to test the protonophoric activity and weak acid properties activity of PZA and POA. **(e)** Dissipation of the pH gradient by POA and CCCP on membranes of pyranine-loaded liposomes formed from mixed POPC, POPG and cholesterol. The black arrow at t_90s_ marks the addition of POA or CCCP. The kinetics of pyranine fluorescence intensity inside liposomes are shown for CCCP (0.4 µg/mL, green curve), CCCP and valinomycin (Val) (0.4 µg/mL and 2.2 µg/L, blue curve), POA (35 µg/mL, red curve), and POA and valinomycin (35 µg/mL and 2.2 µg/L, black curve). After t_600s_, the ionophore Lasalocide A (X537A) was added to the cuvette to equilibrate the pH inside and outside the liposomes. The lipid concentration in the cuvette was 20 µg/mL. **(f)** Dissipation of the pH gradient by PZA or POA on membranes of pyranine-loaded liposomes, the time of addition is marked by the arrow at t_90s_. The kinetics of pyranine fluorescence inside liposomes are shown for POA (140 µg/mL, black curve), PZA (140 µg/mL, red curve), acetate (47.2 µg/mL, blue curve). Grey represents the control condition with vehicle only (DMSO). **(g)** Dose-dependent effect of POA on pH-gradient dissipation on membranes of pyranine-loaded liposomes. Increasing concentrations of POA (35 µg/mL, black curve), (70 µg/mL, red curve), (140 µg/mL, blue curve) were tested. After t_600s_, the control ionophore Lasalocide A (X537A) was added to the cuvette to equilibrate the pH inside and outside the liposomes. The lipid concentration in the cuvette was 20 µg/mL.

Then, we aimed at confirming these results in a system made of pure liposomes which is complementary to the BLM. The cyclic movement of a protonophore through the lipid membrane, which results in a proton transport, is accompanied by a stage of an electrical charge transfer. Hence, in a system of pure liposomes, the generated diffusion potential on the membrane quickly blocks proton transport outside of the vesicles. Consequently, the study of protonophores requires the presence of an electrogenic ion carrier, such as valinomycin which is an electrogenic K^+^ carrier ^63, 64^. So, to confirm our BLM results, we investigated the effect of PZA and POA on the intraliposomal pH-sensitive dye pyranine (Fig 4d) by monitoring its fluorescence intensity profiles in the presence (Fig 4e; black curve) and in the absence (Fig 4e; red curve) of valinomycin. The effect of POA-induced acidification was minor and almost not affected by the addition of valinomycin. In contrast, the effect of the well-established protonophore CCCP was very pronounced in the presence of valinomycin (Fig 4e; blue curve) and almost absent without this electrogenic K^+^ carrier (Fig 4e; green curve). This subset of experiment obtained within the pyranine-loaded liposomes system fully supports our previous BLM results and strongly argues against the protonophoric action of PZA and POA.

Alternatively, we also tested the ability of these compounds to impact the respiratory electron transport chains, which is another well-known characteristic of protonophores. The respiration rate of isolated mitochondria (RLM) was determined by recording the activity of respiratory electron transport chains, which, in turn, is limited by the proton electrochemical gradient. Under the conditions of upper proton motive force limitation, the respiration rate is minimal and depends on the intrinsic proton leakage across the inner mitochondrial membrane. The addition of a protonophore diminishes the proton motive force value and stimulates respiration. In experiments measuring rat liver mitochondrial respiration, both PZA and POA poorly stimulated the respiration, while the addition of 2,4-dinitrophenol (DNP), a known protonophore, drastically stimulated the respiration (Fig S15). This suggests once again that the latter allows the cyclic translocation of protons, while PZA/POA did not.

We capitalized from the pyranine loaded liposome system to test whether PZA/POA have the ability to translocate protons and acidify the bacterial cytosol following a weak acid permeation model. According to this model, the protonation of weak acids into their neutral form increases their ability to cross biological membranes ^65, 66, 67^, thereby raising their concentrations until they reach a steady-state level. More importantly, when protonated, these molecules translocate through the membrane by passive diffusion and carry protons that, based on the pKa of the carboxylic acid function, might be subsequently released ^25, 65^. To test this hypothesis, we treated pyranine-loaded liposomes with PZA or POA (Fig 4f). The well-characterised weak acid acetate, used as positive control, quickly acidified the content of the lumen (Fig 4f; blue curve). If PZA had no effect on intraliposomal pH, POA on the other hand was able to acidify the liposomal lumen (Fig 4f; red and black curves, respectively). Finally, increasing concentration of POA resulted in a dose dependent acidification of the lumen (Fig 4g), therefore fully supporting a conventional weak acid permeation model.

In summary, we propose a unifying model in which acidic pH, and not PanD inhibition, is the primary driver of PZA efficacy against *Mtb* through a weak acid permeation model (Fig 5). In this model, POA behaves as a membrane-permeant weak acid that enters in its protonated form and releases protons intracellularly, collapsing pH homeostasis in a pH-dependent manner. In our experimental system, PanD enzymatic activity and exogenous supplementation with CoA precursors fail to efficiently buffer this acidification process and counteract PZA/POA lethal activity, but confer under less-physiological, such as glycerol-based *in vitro* conditions, an advantage towards PZA/POA action. This provides a molecular mechanistic explanation for why the previously observed PanD-mediated low-level resistance is mainly detectable *in vitro* and rarely identified as a consistent and reliable marker of PZA resistance in clinical setting.

**Figure 5.**
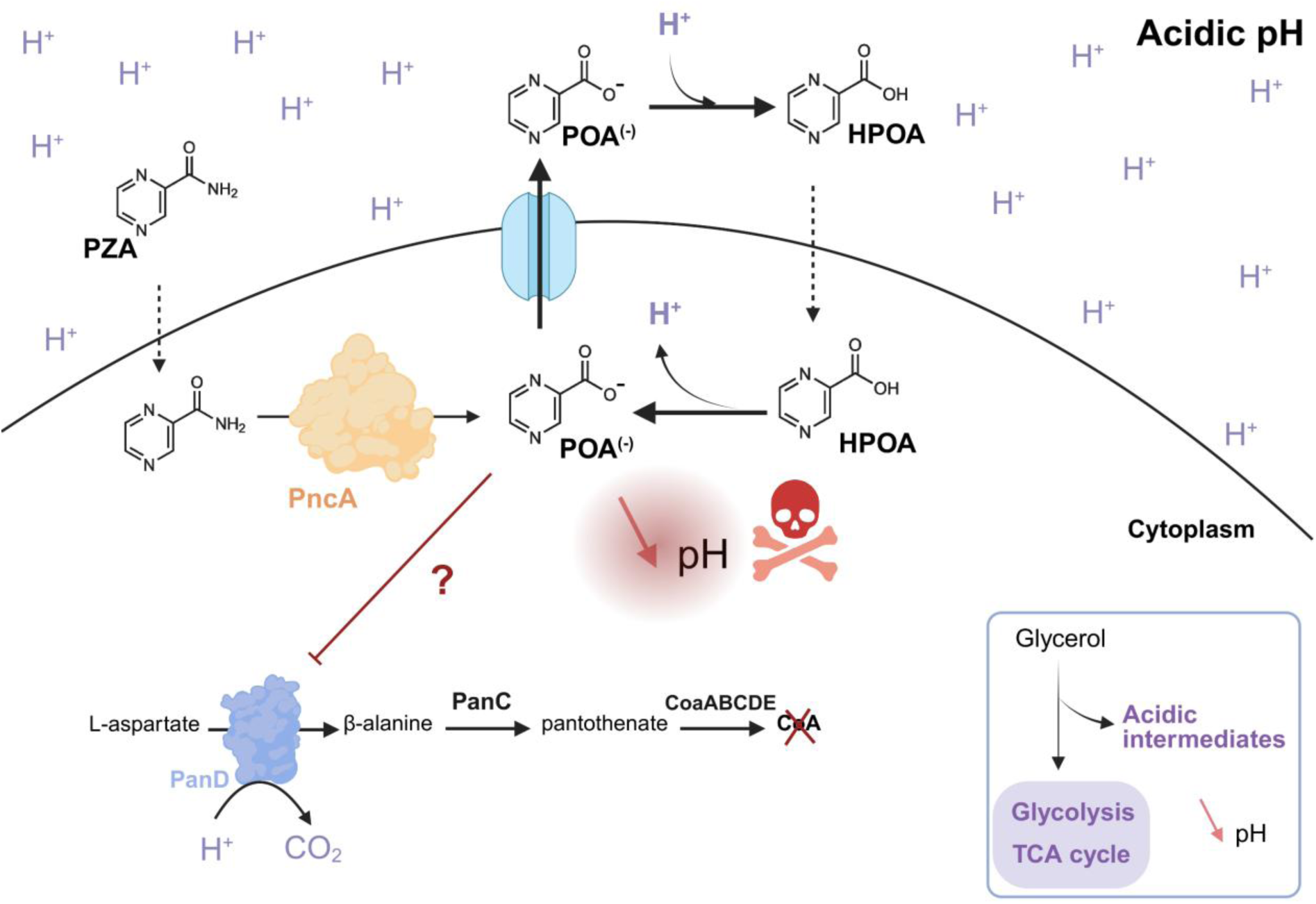
PZA/POA is bactericidal through pH-dependent weak acid permeation and cytosolic acidification as primary mechanism of action. Revisited model of PZA/POA efficacy, supporting a pH-dependent mechanism of weak acid permeation and cytosolic acidification as main mechanism of bactericidal activity. In this model, we propose that the presence of glycerol potentiates PZA/POA efficacy by generating acidic intermediates that lower IBpH. Accordingly, β-ala and Panto might mitigate this effect by acting as buffering molecules that alleviate acidosis. Therefore, in this model PanD minimally contributes to PZA tolerance and cannot be considered as the primary target of the drug.

## Discussion

Despite being used for more than 50 years in clinical settings, the exact mode of action of PZA remains elusive and potentially controversial with two current models biologically conflicted. In this study, we aimed at comprehensively investigate the molecular bases of these two opposite modes of action to decipher the process that underlies PZA efficacy.

By performing antimicrobial susceptibility testing at various pH, we confirmed that acidic pH drastically boosts PZA/POA efficacy *in vitro*. These results are in line with many seminal studies showing that PZA and POA are poorly active in standard Middlebrook culture media ^5^ and strongly potentiated at mildly acidic pH ^23, 24, 25, 68^. These findings are also in agreement with more recent work showing that acidic pH is required for PZA antibacterial efficacy in more complex systems, such as infected macrophages ^30, 69^ or lungs ^48, 70, 71^. By using high-resolution correlative nano-secondary ion mass spectrometry imaging and scanning electron microscopy, we indeed previously demonstrated that a functional endolysosomal network displaying acidic features greatly increases PZA/POA enrichment accumulation in *Mtb* or its surrounding vicinity. Pharmacological or genetic modulation of *Mtb* exposure to such acidic environments had also direct impact on PZA/POA enrichment at the bacterial single-cell level, and ultimately a potent antibacterial activity ^69^.

Capitalizing from the well-established *Mtb* pH-GFP reporter system ^29, 30, 43, 46, 47^, we set up distinct *in vitro* assays to monitor non-invasively PZA/POA effect on *Mtb* IBpH. Across all formats, both compounds lowered the IBpH of *Mtb* cytosol in a dose- and pH-dependent manner. This specific feature was also observed previously with the reference strain H37Rv during host-cell infection, where prolonged exposure to higher drug concentration leads to IBpH acidification ^30^. The use concanamycin A, a conventional v-ATPase inhibitor that impairs protons import within endolysosomes, further completely abolished PZA-mediated effect on IBpH and abrogated PZA antimicrobial activity ^30, 69^. This work showed that functional endolysosomal acidification is also a critical requirement for PZA/POA pH-disruptive property within host-cells.

In the current study, we show that PZA/POA-driven IBpH acidification targets both actively growing and non-replicating *Mtb* cells demonstrating that PZA/POA antimicrobial activity does not rely on *Mtb* replication unlike other anti-TB drugs such as INH ^72, 73^. This feature observed onto non-replicating cells is in accordance with previous reports from Darby *et al*. and Fontes *et al*. who relied on genetically-encoded or probe-based IBpH reporters to demonstrate that PZA/POA display pH-homeostasis disruptive activity in buffers ^28, 29^. Moreover, our data show that even if the abundance of a given carbon source, such as glycerol, can alter PZA/POA efficacy, *Mtb* killing by PZA/POA does not require the presence of a growth permissive carbon source.

Altogether, our data support a direct correlation between extracellular protons availability, PZA/POA-induced IBpH collapse and growth inhibition providing strong experimental evidences for the pH-dependent PZA-mediated cytosolic acidification model originally proposed by Zhang *et al.* ^24, 26^.

To test whether the marked IBpH alteration observed in phosphate citrate buffer leads to bacterial killing, we performed time-, dose- and pH-dependent killing experiments. In phosphate citrate buffer adjusted at pH5.5, PZA displays strong bactericidal activity. Indeed, a 14-day exposure to 200 or 500 μg/mL of PZA in buffer adjusted at pH 5.5 led to a two to three-fold Log^10^ reduction in bacterial viability, whereas no killing was observed at pH 7. This result echoes previous studies showing pH-dependent killing on non-replicating bacteria at acidic pH ^60^, yet differs from Peterson *et al*. who reported a two Log^10^ reduction in bacterial viability upon treatment with 200 μg/mL of PZA in PBS buffer at pH 7 ^35^. This difference in killing efficacy at neutral pH could be due to distinct inoculum density, or alternatively the weaker buffering capacity of PBS in comparison to phosphate citrate buffer, which could facilitate local acidification during prolonged incubation. Despite these discrepancies, the collective evidences support a model in which PZA rather acts as a slow-killing drug in growing or non-growing conditions *in vitro*; a conclusion recently reaffirmed by Gouzy *et al*. using a custom culture media ^58,74^.

Our current work also shows that structurally related analogues to PZA/POA, display similar pH-dependent behaviour in term of IBpH homeostasis perturbation and antibacterial activity. This suggests that such mechanism is not unique to PZA/POA but rather conserved among structurally related molecules. Analysis of the chemical structure of this sub-class of compounds clearly supports a model in which harbouring a carboxylic acid function, or an amide readily convertible into a carboxylic acid function though enzymatic action, accounts for this phenotype, consistent with a weak acid permeation model ^43, 44^.

A long-standing debate regarding the pH-dependent model of PZA/POA action is whether the proton shuttling effect and the resulting IBpH acidification are a direct effect of the drug or the consequence of a yet-unknown inhibited process ^40^. To resolve this, we designed an *in vitro* assay, using phospholipid-based liposomes, providing a minimal system devoid of proteins or metabolic pathways. We show that POA, the bio-active form of the drug, is able to lower the luminal pH of these liposomes in a pH-dependent manner. The most parsimonious explanation to this process, is that HPOA alone transports proton inward, in strict accordance with the Henderson–Hasselback relationship. These findings demonstrate that POA-driven acidification does not depend on inhibiting any cellular process; rather, it reflects an intrinsic, pH-dependent proton-translocation mechanism.

Because compound-mediated IBpH acidification can arise by two distinct processes (*e.g*. protonophore or membrane permeant weak acid), we aimed at determining the molecular basis of this acidification process. Recent studies have suggested that the PZA/POA pair might act as protonophore that flips protons across the mycobacterial envelope. In this study, we show by using three independent experimental systems, the BLM, the RLM and fluorescein-loaded liposomes that PZA/POA is not a protonophore *per se*. In contrast to well-characterized molecule CCCP, PZA/POA don’t repeatedly flip protons across biological membranes ^39, 40, 75^. Instead, we provide strong evidence that POA acts as weak acid that diffuses across the membrane under its neutral protonated form, and release proton upon entry within the liposomal lumen until reaching an equilibrium. These results, together with previous observations that the acid-base equilibrium of POA^-^/POA is critical for efficacy ^25, 28^, demonstrate that PZA/POA acidifies IBpH through a weak-acid permeation mechanism rather than by protonophore-mediated proton shuttling.

Recently, through an integrative approach relying on the use of the custom HBO medium and combining genome-wide CRISPRi profiling, metabolomic analyses, and comparative genomics, Gouzy *et al.* discovered that the ion channel Rv2571c facilitates the efflux of α-ketoglutarate and sensitizes *Mtb* against PZA ^74^. Interestingly, release of α-ketoglutarate exacerbates PZA-induced IBpH acidification under acidic conditions, characteristic of the host environment ^74^. Loss-of-function mutations in Rv2571c conferred PZA resistance both *in vitro* and *in vivo*, underscoring the physiological relevance of this new pathway and solidifying cytoplasmic acidification as the central driver of PZA’s bactericidal activity ^74^.

Finally, another key biological question addressed in our study, is the potential direct contribution of the aspartate decarboxylase PanD in PZA/POA efficacy. Antimicrobial susceptibility assays in standard 7H9 broth (circumneutral or acidic) showed that supplementation with Panto or β-ala, strongly antagonised PZA/POA efficacy, as previously reported ^31, 32, 45^. We also confirmed that the nature of the carbon source is a key parameter in efficacy, with glycerol having potentiating effect on this drug ^38, 76^. Following this observation, we tested whether the observed antagonisms were carbon-source dependent. Intriguingly, while glycerol was replaced by oleic acid as the sole carbon source ^58^, Panto-mediated antagonism was almost completely abolished. Exploiting this observation, we compared the PZA/POA susceptibility of the *panCD*-deleted auxotroph H37Rv mc^2^ 6206 ^77^ with prototrophic H37Rv parent. In the oleic acid-based medium (HBO medium), results highlight that the lack of functional PanCD does not affect PZA/POA efficacy, suggesting that PanD is unlikely to be the primary target of the drug. These data are also in accordance with the IBpH homeostasis assays that revealed that both prototrophic and auxotrophic strains are affected by PZA/POA-mediated acidification and that exogenous Panto or β-ala supplementation did not revert this phenotype. This IBpH homeostasis alteration was also observed on non-replicating starved cells in phosphate citrate buffer in a pH-dependent manner, and time-kill experiments showed that deletion of the *panCD* genetic locus did not alter bacterial viability even after 14 days of deprivation, as previously reported ^62^. On the other end, PZA treatment triggered a slow but very strong decrease in bacterial viability regardless of the presence of PanCD enzymes or the exogenous supplementation of Panto. Collectively, these data argue that PZA/POA act through a PanD-independent, pH-dependent protons-import mechanism.

Of note, while performing these experiments we observed some minor differences between a strain harbouring a functional PanD in comparison to a PanD-deficient strain. Indeed, in several assays the PanD-deficient strain displayed a slightly greater buffering deficit after PZA/POA exposure. Because PanD convert L-Asp to β-ala while consuming a proton, we initially proposed that active PanD can buffer cytosolic pH under low-level drug or stress. Overexpressing PanD from *Mtb*, *M. smegmatis* or *E. coli* indeed confers low-level resistance to POA on standard 7H10 media, and *panD* or *clpCP* mutations that stabilise PanD were isolated in low-grade POA-resistant mutants ^31, 32, 33^. To support this model, we overexpressed *panD* and performed IBpH monitoring in the presence of increasing PZA concentrations with or without precursors of the CoA biosynthesis pathway. Our data showed no major differences in between the *Mtb* WT or overexpression strain suggesting that PanD activity alone, unlike mitigates mild PZA/POA-mediated acidification.

Another, interesting feature that could bridge the two models is that β-ala and Panto are known buffers that alleviate metabolic acidosis in eucaryotic cell ^78, 79^. Their partial protection in *Mycobacterium* towards cytosolic acidification might arise from the same physicochemical buffering rather than pathway rescue, a hypothesis that warrants deeper investigations.

It is also worth mentioning that over the past few years, several putative targets have been proposed to be involved in PZA/POA susceptibility therefore leading to the concept that PZA/POA might act as a multi-target inhibitor ^39, 40, 41, 42^. Nevertheless, while this poly-pharmacology hypothesis remains to be validated, our current data, alongside recent genome-wide CRISPRi ^74^ screens and clinical genomics ^80^, strongly supports the pH-dependent model as the dominant mechanism underlying its bactericidal activity. Indeed, proposed PZA targets such as PanD and GpsI were not identified as PZA-sensitive hits in a recent CRISPRi screen conducted under acidic conditions, where PZA exhibits high activity. Furthermore, neither gene show evidence of positive selection in clinical isolates, as would be expected for a bona fide drug target, such as *rpoB*, the well-established target of rifampicin.

Finally, our study underscore how profoundly experimental context shapes PZA efficacy. The artificially high concentration of glycerol has long been known to modulate antibiotic susceptibility ^55, 56, 76, 81, 82^. Yet, to our knowledge, it is the first instance in which a pronounced glycerol-driven antagonism, has been interpreted as evidence for an accepted anti-TB drug mode of action that appears limited to *in vitro* conditions. In our case, assays conducted in the more physiological, acidic HBO medium, revealed PZA’s full antibacterial potency ^74^, highlighting its status as an unusual drug whose activity is maximised under conditions mimicking better the environment *Mtb* experiences during infection.

Taken together, our data support a revised model (Fig 5) in which acidic pH serves as the principal factor of PZA/POA efficacy against *Mtb.* In this framework, POA behaves as a membrane-permeant weak acid. Our findings also indicate that PanD is unlikely to be the primary target of PZA. Its decarboxylase activity, along with the buffering capacity provided by β-ala and Panto, poorly counteract PZA/POA-driven acidification and bacterial killing. While a strong antagonism was reported, it was only observed under certain *in-vitro* conditions. Notably, this protective effect is abolished under more physiological settings such as the acidic HBO medium, explaining why PanD-associated low-level resistance is rarely seen in clinical isolates and offers limited value as a reliable marker of PZA resistance.

## Materials and Methods

### Bacterial strains and culture conditions

Laboratory bacterial strains *Mtb* H37Ra ATCC 25177, *Mtb* H37Rv WT, *Mtb* H37Rv mc^2^ 6206, *Mtb* H37Rv mc^2^ 6230, *Mtb* H37Rv Δ*pncA* and *Mtb* H37Rv Δ*pncA::pncA* were used in this study. *Mtb* H37Rv mc^2^ 6206 (Δ*panCD*Δ*leuCD*) and mc^2^ 6230 (Δ*panCD*Δ*RD1*) strains ^51, 52, 77^ were kindly provided by William R. Jacobs, Jr. (Albert Einstein College of Medicine, New York, NY, USA). The mutant Δ*pncA* as well as its related WT were gifted by Helena I. M. Boshoff (NIH, Bethesda, USA) as previously described ^83^.

Bacterial cultures were routinely grown on standard Middlebrook 7H9 broth (BD Difco; #271310) supplemented with 0.2% glycerol (*v/v*) (Euromedex; #EU3550), 0.025% (*v/v*) tyloxapol (Sigma-Aldrich; #T8761) and 10% (*v/v*) oleic acid, albumin, dextrose, catalase (OADC enrichment - BD Difco; #211886). When required, Panto (Sigma-Aldrich; #21210) was used at a final concentration of 25 μg/mL to support the growth of *Mtb* H37Rv mc^2^6230 and mc^2^6206, and 50 μg/mL leucine (Sigma-Aldrich; #L8000) and 2g/L casamino acids (VWR; #J851) were also used to support the growth of *Mtb* H37Rv mc^2^6206. Recombinant *Mtb* H37Ra ATCC 25177 and *Mtb* mc²6230 strains expressing pH-GFP ^46^ (pUV15-pHGFP; Addgene plasmid #70045, kindly gifted by Sabine Ehrt), were generated by electroporation according to Goude *et al*. ^84^ and subsequently selected onto 7H11 agar medium (BD Difco; #283810) supplemented with 10% OADC and 50 μg/mL hygromycin B (Toku-E; #H007). Hygromycin B was used as selection marker for the culture maintenance of the fluorescent strain at a final concentration of 50 μg/mL, but was not used during IBpH homeostasis disruption experiments.

### Generation and validation of *Mtb* overexpression and complementation strains

To construct a *panCD* overexpression strain, a genomic fragment encompassing the adjacent genes *panC* and *panD* was amplified by PCR from *Mtb* H37Ra genomic DNA using primers #P1 5’ TATATTAATTAAATGACGATTCCTGCGTTCCATCC 3’ and #P2 5’ TATAAAGCTTCTAGTGGTGGTGGTGGTGGTGTCCCACACCGAGCCGGGG 3’. DNA fragment was purified (NucleoSpin® Gel and PCR Clean-up, Macherey-Nagel) and further digested with PacI and HindIII restriction enzymes (New England Biolabs, #R0547L and #R3104L). Simultaneously, pUV15-pHGFP was digested with PacI and HindIII to remove the pH-GFP coding sequence. Purified and digested insert and vector were then ligated using T4 DNA ligase (New England Biolabs; #M0202L) to generate the pMSD42 vector. Subsequently, pMSD42 vector was digested with SpeI (New England Biolabs, #R3133L) and HindIII restriction enzymes. The 1758 bp insert was purified and cloned at the XbaI (New England Biolabs, #R0145L) and HindIII sites of the pMV306 ^53, 85^ digested vector to generate the pMSD44 vector. Accordingly, pMSD42 (hygromycin resistant, episomal) and pMSD44 (kanamycin resistant, L5-integrative) both harbor the *panCD* locus in frame with a C-terminal 6xHis-tag under the control of the Psmyc promoter. Empty pMV306 and pMSD44 were subsequently used to electroporate *Mtb* H37Ra pH-GFP. PanD production was further confirmed by immunoblot. Briefly, recombinant *Mtb* pH-GFP harboring pMV306 and pMSD44 were grown in complete 7H9 medium supplemented with 50 µg/mL of hygromycin and kanamycin, and incubated at 37°C under shaking until OD_600nm_ reached late exponentially growing phase (approximately OD_600nm_ 1.0-2.0). Samples were harvested for 10 min at 3,500rpm, then washed twice with PBS (pH 7.4) buffer containing 0.05% (v/v) Tween-20. All samples were re-suspended with PBS (pH 7.4) buffer for normalization based on OD_600nm_. Then, bacterial cells were mixed with 200 μL of 0.1 mm diameter glass beads (BioSpec), and mechanically disrupted during 3 × 4 min of vigorous shaking using Mini-Beadbeater-96 (BioSpec, USA). The lysates were centrifuged for 5 min at 5,000 rpm and the resulting supernatant (∼750 µL) was carefully collected. All conditions were normalized a second time by Bradford based on protein content. Samples were diluted in Laemmli Sample Buffer, incubated at 95°C for 5-10 min and further stored at -20°C. Each sample was then loaded and separated on to 15% SDS/PAGE and further transferred on to a 0.2 µm PVDF membrane (Bio-Rad, #1704156) using a Trans-Blot Turbo Transfer System (Bio-Rad). Immunoblotting of 6×His-tagged proteins was performed using anti-HisProbe™ HRP conjugate according to the manufacturer’s instructions (Thermo Fisher Scientific, #15165). Coomassie blue/Ponceau Red Staining and/or detection of GroEL1 levels through its natural poly-histidine sequence present in its C-terminal domain were used as internal loading controls. Detection was achieved using Pierce™ ECL Western Blotting substrate solution (Thermo Fisher Scientific, #32132) and visualized with a ChemiDoc™ MP Imaging System (Bio-Rad). The complemented strain Δ*pncA::pncA* was constructed by integrating a plasmid (attL5 site) allowing the expression of the *pncA* gene under its native promoter (300bp upstream of start codon). Complementation of the *Mtb* H37Rv mc^2^ 6206 (Δ*panCDΔleuCD*) strain with *panCD* was done by integrating a plasmid (attL5 site) allowing the expression of the *panCD* operon under the *hsp60* promoter.

### Antimycobacterial susceptibility testing and MIC determination

MIC determinations against *Mtb* H37Ra ATCC 25177 were performed at circumneutral and mildly acidic pH as previously described ^43^. Briefly, all experiments at pH 6.8 were performed in standard Middlebrook 7H9 medium containing 0.2% glycerol and 10% OADC enrichment. To avoid Tween-80-mediated toxicity, 0.025% Tyloxapol (Sigma-Aldrich; #T8761) was used. For pH-dependent experiments, the medium was supplemented with 50 mM MES and adjusted to pH 5.5 ^86, 87^.

Chemical compounds including PZA (#P7136), POA (#P56100), NAM (#N0636), NAC (#N4126), SA (#247588), INH (#I3377), ETH (#E6005), PTH (#SMB00387), RIF (#R3501) were purchased from Sigma-Aldrich (Saint-Quentin Fallavier, France). All compounds were > 95% purity. Kanamycin (KAN) sulfate (#UK0010D) was purchased from Euromedex. Compounds were dissolved in dimethyl sulfoxide (DMSO) (DASIT group; #455103) at 50 mg/mL for PZA, POA, NAM, NAC, and KAN, at 20 mg/mL for SA, at 10 mg/mL for INH, at 2 mg/mL for ETH, PTH and RIF. The solutions were then sterilized using 0.22 µm syringe filters (#146560) from Clearline.

Two-fold serial dilutions of the compound stock solutions were used to obtain a range of 10-12 concentrations in the appropriate culture medium. Exponential-phase *Mtb* cultures (OD_600nm_ that approximates 0.6-1.2) were adjusted to a bacterial density of 5 × 10^6^ CFU/mL in complete 7H9 medium at pH 6.8 or pH 5.5. A volume of 100 μL of this inoculum was then dispensed into each well containing 100 μL of each serial dilution of the compound, achieving a final bacterial concentration of 2.5 × 10^6^ CFU/mL per well. Positive growth controls (*i.e.*, inoculum without antibiotics or with DMSO), negative growth controls (*i.e.*, 50 μg/mL KAN), and sterility controls (*i.e.*, medium only) were included. The 96-well flat-bottom Nunclon Delta Surface microplates with lid (Thermo-Fisher Scientific; #167008) were incubated at 37 °C for 14-21 days for both pH conditions. The minimum inhibitory concentration (MIC) of each antibiotic at pH 6.8 or pH 5.5 was visually determined in accordance with the EUCAST and CLSI guidelines ^88^. Results were confirmed by OD_600nm_ measurement using the TECAN spark 10M™ multimode microplate reader. The lowest antibiotic concentration that inhibited bacterial growth was determined as the MIC which almost perfectly aligned with the MIC_90_ calculated from OD_600nm_ measurement.

Antagonistic studies were performed in the exact same manner in the presence of Panto which was dissolved in water at 25 mg/mL and used at a final concentration of 25 µg/mL. This concentration was chosen and conserved throughout the study as it is known to fully restore the growth of an *Mtb* auxotrophic strain ^51^. For visual representation of potentiating and antagonistic effects, MIC have been expressed as Log^2^ Fold-Change (MIC) in comparison to their respective control condition. The corresponding heat-map has been generated accordingly.

Finally, when required, a half-maximal effective concentration (EC_50_) was determined using a four-parameter log-logistic nonlinear regression model (R Studio software, The R Project for Statistical Computing), as the concentration required to achieve 50% of the observed effect (i.e., *Mtb* growth inhibition). All the presented data have been obtained from experiments performed in technical triplicate at least on three biological replicates.

### Antimycobacterial susceptibility testing in High-BSA-Oleate (HBO) media

To prepare High-BSA-Oleate (HBO) media, Middlebrook 7H9 medium was supplemented with 50 g/L of fatty acid-free bovine serum albumin (BSA) (Millipore; #3117057001), 3mM oleic acid (Millipore-Sigma; OX-0165), 0.850g/L NaCl, tyloxapol 0.05% (Sigma-Aldrich; #T8761;) and 50mM of MES buffer (Sigma-Aldrich; #M3671). Glycerol was omitted. The media was acidified to pH 5.8 by the addition of concentrated HCl. A stock of PZA at 100 mg/mL in DMSO was used to distribute a gradient concentration of PZA in 384 well-plates using a HP dispenser. Cultures of mid-log phase bacteria (OD_600nm_ = 0.5-0.8) were washed three times in PBS and inoculated in HBO media pH5.8 to a final OD_600nm_ of 0.05. A volume of 50 µL of bacteria were added per well. Plates were incubated for 14 days at 37°C before OD_600nm_ reading using a plate reader. In the case of *Mtb* H37Rv mc^2^6206, the media were supplemented with Panto (25 µg/mL), leucine (50 µg/mL) and casamino acids (2 g/L) to support bacterial growth.

### Intrabacterial pH homeostasis disruption assays

Recombinant *Mtb* ATCC 25177 and *Mtb* mc²6230 strains expressing the ratiometric pH-sensitive pH-GFP sensor were used to monitor IBpH homeostasis and drug-mediated perturbations on both exponentially growing and non-replicating *Mtb* cells ^29, 30, 43, 46^.

*Mtb* IBpH determination on exponentially growing *Mtb* was performed as previously reported ^29, 30, 43, 46^. Briefly, exponentially growing *Mtb* cultures (OD_600nm_ 0.5-1.5) were centrifuged at 3,500 rpm for 10 min. The pellet was resuspended in 10 mL of Middlebrook 7H9 medium adjusted at pH 5.5 or pH 6.8 in order to have a normalized OD_600nm_ of 0.8. Then, 100 µL of this bacterial inoculum was used to inoculate wells containing 100 µL of serially diluted compounds of interest, resulting in a final OD_600nm_ of 0.4 in each well. When required the medium was supplemented with Panto at a final concentration of 25 μg/mL. After 24 h of incubation, the fluorescent pH-GFP signal was subsequently detected using excitation/emission wavelengths of λ_ex405nm_/λ_em535nm_ followed by λ_ex488nm_/λ_em535nm_ using a TECAN spark 10M™ multimode microplate reader (Tecan Group Ltd, France). Then fluorescence intensity ratios were calculated from data obtained at λ_em535nm_ from λ_ex405nm_/λ_ex488nm_ excitations. For experiments carried at pH 6.8, the well-established protonophore Carbonyl Cyanide *m*-Chlorophenylhydrazone (CCCP) (Sigma-Aldrich; #215911) was used as positive control of IBpH homeostasis disruption; while for experiments conducted at pH 5.5, the well-established ionophore Monensin (MON) (Sigma-Aldrich; #M5273) was used as positive control.

*Mtb* IBpH determination on non-replicating *Mtb* was performed as described above with slight modifications. Approximately, 50 mL of exponentially growing *Mtb* cultures (OD_600nm_ 0.5-1.5) were split into 5 independent 15 mL conical tubes and centrifuged at 3,500 rpm for 10 min. The pellets were washed once and re-suspended in 10 mL of phosphate citrate buffer with pH ranging from 7 to 5.5 by increments of 0.5 to have a normalized OD_600nm_ of 0.8. Then, 100 µL of this bacterial inoculum was used to inoculate wells containing 100 µL of serially diluted compounds of interest, resulting in a final OD_600nm_ of 0.4 in each well. After 48 h of incubation, the fluorescent pH-GFP signal was recorded and expressed as fluorescent ratio as described above.

To convert pH-GFP ratios into pH units and determine *Mtb* IBpH, a calibration curve was performed. Recombinant *Mtb* H37Ra ATCC25177 expressing pH-GFP was cultivated in standard 7H9 culture medium supplemented with 10% OADC enrichment and 50 μg/mL hygromycin B until exponential phase (OD_600nm_ between 0.5-1.5). Six independent conical 15 mL-tubes containing 10 mL of bacterial culture were prepared and centrifuged at 3,500 rpm for 10 min. The supernatant was discarded, and the bacterial pellet was re-suspended in 1 mL of phosphate citrate buffer with pH values ranging from 8 to 5.5 in 0.5-unit increments. Following a 2-hours incubation at 37 °C, the bacterial cells were mixed with 200 μL of 0.1 mm diameter glass beads (BioSpec), and disrupted during 3 × 4 min of violent shaking using Mini-Beadbeater-96 (BioSpec, Bartlesville, OK, USA). The lysates were centrifuged for 5 min at 1,500 rpm, the resulting supernatant (∼750 µL) was carefully collected, and 3 × 200 µL were transferred into each well of a 96-well plate (Nunclon Delta Surface microplate). pH-GFP signal was detected using excitation/emission wavelengths of λ_ex405nm_/λ_em535nm_ followed by λ_ex488nm_/λ_em535nm_ using the TECAN spark 10M™ multimode microplate reader. Fluorescent ratios were used to build a non-parametric calibration curve, by applying the locally estimated scatterplot smoothing (LOESS) prediction model in R (R Studio software, The R Project for Statistical Computing, version 1.3.1073) that fits the obtained data to a smooth standard curve. This LOESS prediction model was then used to correlate fluorescence ratio with their respective IBpH values. All the results were exported as CSV files, imported into the R Studio software, and graphical representations were plotted using the ggplot2 package (version 3.5.0). Results of all the presented data have been performed in technical triplicate at least on three biological replicates.

For experiments with mc^2^6206 (Δ*panCD*Δ*leuCD*) and Δ*pncA*, as well as their wildtype and complemented counterparts, IBpH homeostasis disruption assays were done in biosafety level 3 laboratory following a similar protocol and suing a Molecular Devices M5 SpectraMax plate reader.

### PZA bactericidal activity on non-replicating *Mtb* using spot CFU-based assays

Exponentially growing cultures of *Mtb* H37Ra pH-GFP and *Mtb* mc² 6230 pH-GFP were centrifuged, the supernatant was removed, the pellets were washed and resuspended in phosphate citrate buffer adjusted to pH 7 or pH 5.5, and supplemented with 50 μg/mL hygromycin B to obtain a final OD_600nm_ of 0.1. PZA was used at final concentration of 0, 50, 200 or 500 μg/mL. MON or RIF compound was used as internal positive growth inhibition control. When required, 25 μg/mL of Panto was also added to the medium. At each time point (day 0, 2, 7 and 14), 200 μL of the undiluted condition (10^0^) was sampled. From this condition, serial dilutions were carried out to obtain the dilution ranging from 10^-1^ to 10^-5^. Then, 10 μL of each dilution were spotted onto a 7H10 agar plate supplemented with 10% OADC, 50 μg/mL hygromycin B and 25 μg/mL Panto to recover *Mtb* cells. The plates were incubated 21 days at 37 °C. Scanning of the plates was performed using a ChemiDoc™ MP Imaging System (Bio-Rad) where bacterial spots imaging was achieved by using the ‘White Epi Illumination’ setting combined with the ‘Standard Filter’. Experiments were performed on two independent occasions and are representative of one replicate.

### PZA bactericidal activity on non-replicating *Mtb* using quantitative CFU analysis

For survival assays in phosphate citrate buffer at pH7 and pH5.5, bacterial cultures were grown until mid-log phase, washed three time in PBS-tyloxapol 0.05% and started at an OD_580nm_ of 0.1 in the absence of drug (CTRL) or in the presence of PZA at 200 or 500 µg/mL. At day 0, 7 and 14, cultures were serially diluted in PBS-tyloxapol 0.05% and plated on 7H10 agar containing 0.5% glycerol and supplemented with OADC and 4 g/L activated charcoal. Plates were incubated at 37 °C for 3 weeks prior to CFU enumeration. All strains tested (WT, mc^2^6206 (Δ*panCD*Δ*leuCD*) and complemented strain Δ*panCDΔleuCD*::*panCD*) were supplemented with 50 μg/mL leucine and 2 g/L casamino acids during the 14-days period. Panto at 25 μg/mL was supplemented to all strains, or not, to evaluate its role in Mtb survival in non-replicating conditions.

### Isolation of rat liver mitochondria

Rat liver mitochondria were isolated by differential centrifugation ^89^ in a medium containing 250 mM sucrose, 5 mM MOPS, 1 mM EGTA, pH 7.4. The final washing was performed in the medium containing bovine serum albumin at final concentration 0.1 mg/mL. Protein concentration was quantified using the Biuret method. All animal handling and experimental procedures were conducted in accordance to the international guidelines for animal care and use and received approval from the Institutional Ethics Committee of the A.N. Belozersky Institute of Physico-Chemical Biology at the Moscow State University (Protocol #3 on February 12, 2018).

### Mitochondria respiration

Respiration of isolated mitochondria was measured using a standard polarographic technique with a Clark-type oxygen electrode (Strathkelvin Instruments, UK) at 25°C using the 782 Oxygen System software. The incubation medium contained 250 mM sucrose, 5 mM MOPS, and 1 mM EGTA, pH 7.4. The mitochondrial protein concentration was 0.8 mg/mL. The reported oxygen uptake values are expressed as nmol/min/mg of protein.

### Membrane potential measurement in isolated mitochondria

Mitochondrial membrane potential was estimated using safranine O dye ^90^. Fluorescence was monitored with a Panorama Fluorat 02 spectrofluorimeter (Lumex, Saint-Petersburg) at λ_ex520nm_/λ_em580nm_. At the end of each measurement, 200 nM CCCP was introduced to collapse the membrane potential. The incubation medium consisted of 250 mM sucrose, 5 mM MOPS, 0.5 mM KH_2_PO_4_, 1 mM EGTA, 2 μM rotenone, 5 mM succinate (pH 7.4), 1 μg/mL oligomycin, 15 μM safranine O. The mitochondrial protein content ranged from 0.6 to 0.9 mg protein/mL and the temperature was maintained constant at 26 °C.

### Planar bilayers

Bilayer lipid membrane (BLM) was created using the brush technique ^91^ from a 2% decane solution of phosphatidylcholine derivative, diphytanoylphosphatidylcholine (DPhPC) (Avanti Polar Lipids; Alabaster, AL). on a 0.6-mm aperture in a Teflon septum separating the experimental chamber into two compartments of equal 3-ml volume. The electrical current across the BLM was measured under voltage-clamp conditions with two AgCl electrodes placed into the solutions on the two sides of the BLM through agar bridges, using a Keithley 428 amplifier (Cleveland, Ohio, USA). The voltage applied to the BLM was set at 50 mV.

### Pyranine-loaded liposomes

The pH of liposomes was measured as previously described by Chen *et al*. ^64^ with slight modifications. To prepare pyranine-loaded liposomes, a lipid mixture from POPC, POPG and cholesterol (5.4, 1.5 and 3.1 mg, respectively) dissolved in a chloroform suspension was dried in a round-bottom flask under a nitrogen stream. The lipid was then resuspended in 1 mL buffer (100 mM KCl, 20 mM MES, 20 mM MOPS, 20 mM Tricine titrated with KOH to pH 8.0) containing 0.5 mM pyranine. The suspension was vortexed and freeze-thawed three times. Unilamellar liposomes were then prepared by extrusion through 0.1 µm pores (Nucleopore polycarbonate membranes) using an Avanti Mini-Extruder. Untrapped pyranine was removed by passage through a Sephadex G-50 coarse column equilibrated with the same buffer solution. The liposomes were diluted in the same buffer at pH 6.0 and final lipid concentration in the cuvette of approximately 20 μg/mL. To measure the pH dissipation rate in liposomes, the suspension was supplemented with 2 mM *p*-xylene-*bis*-pyridinium bromide to quench the fluorescence of leaked pyranine. The internal pH was estimated by measuring the fluorescence intensity at λ_ex455nm_/λ_em505nm_, as the fluorescence emission of liposome-loaded pyranine monotonously increases upon excitation at λ_ex455nm_ over a pH range of 6.0 to 7.9 ^64^. The fluorescence was monitored on a Panorama Fluorat 02 spectrofluorimeter (Lumex, Saint-Petersburg) ^92^. At the end of each recording, 1 µM X537A (Lasalocid A) was added to dissipate the remaining pH gradient. In several experiments, to prevent the formation of H^+^-diffusion potential, the experiments were conducted in the presence of 10 nM K^+^-carrier valinomycin. To minimize the leakage of H^+^ from liposomes, the experiments were performed at 15°C.

### Quantification and statistical analysis

All results displayed in this study were obtained from biologically independent experiments (*n* = 3) each performed in at least three technical replicates unless otherwise stated. Statistical analysis for IBpH homeostasis disruption assays was performed comparing the means between the conditions of interest using one-way ANOVA followed by Tukey’s post-test with the ‘aov()’ and ‘TukeyHSD()’ functions in R Studio using the ggpubr R package (The R Project for Statistical Computing). All *p*-values presented in the text or figures are relative to the control condition or to the lowest drug concentration applied in dose-response experiments.

Statistical analysis for quantitative CFU assays was performed by comparing the means between the conditions of interest in R Studio (The R Project for Statistical Computing). Briefly, the ‘var.test()’ function to assess variance homogeneity in between two groups of interest. Since variances were equal (*p* > 0.05), a Student Two Sample t-test was applied using the ‘t.test(…, var.equal = TRUE)’ function. In the main manuscript and supplementary material exact *p*-values were added to the figures to enable readers to assess statistical significance. Non -statistical significance was considered for *p*-value ≥ 0.05 and abbreviated as ‘n.s’. Each figure or table legend specifies the statistical tests used, the number of biologically independent replicates and the number of technical replicates.

## Abbreviations

PZA: Pyrazinamide
TB: Tuberculosis
Mtb: Mycobacterium tuberculosis
POA: Pyrazinoic acid
INH: Isoniazid
RIF: Rifampicin
EMB: Ethambutol
NAD+: Nicotinamide adenine dinucleotide
FAS I: Fatty acid synthase I
CoA: Coenzyme A
MIC: Minimal inhibitory concentration
NAM: Nicotinamide
NAC: Nicotinic acid
SA: Salicylic acid
ETH: Ethionamide
PTH: Prothionamide
β-ala: β-alanine
L-Asp: L-aspartate
Panto: Pantothenate
MON: Monensin
CCCP: carbonyl cyanide *m*-chlorophenyl hydrazone
IBpH: Intrabacterial pH
LOESS: Locally estimated scatterplot smoothing
EC_50_: Half maximal effective concentration
HBO: High-BSA-Oleate
BLM: Bilayer lipid membrane
DNP: 2,4-dinitrophenol

## Acknowledgments

We would like to acknowledge all members of the Lipolysis and Bacterial Pathogenicity group and the LISM unit for continuous support and insightful discussions. Special thanks are addressed to Maximiliano Gutierrez, Ruben Hartkoorn, Julien Vaubourgeix, Benjamin Ezraty, Julie Viala and Bérengère Ize for fruitful discussions. We also would like to thank Vadim Makarov for providing the PZA/POA compounds to perform the bio-electrophysiology experiments.

## Authors contribution

**Conceptualization** JL, ARB, YNA, AG, PS

**Formal Analysis** JL, YNA, AG, PS

**Funding Acquisition** YNA, AG, PS

**Investigation** JL, TIR, AA, VP, AMF, LSK, JFC, MSD

**Methodology** JL, ARB, YNA, AG, PS

**Project Administration** PS **Supervision** SC, GG, YNA, PS

**Visualization** JL, YNA, AG, PS

**Writing – Original Draft Preparation** JL, YNA, AG, PS

**Writing – Review & Editing** JL, TIR, AA, VP, AMF, LSK, MSD, GG, JFC, SC, ARB, YNA, AG, PS

## Supporting information

**Figure S1.**
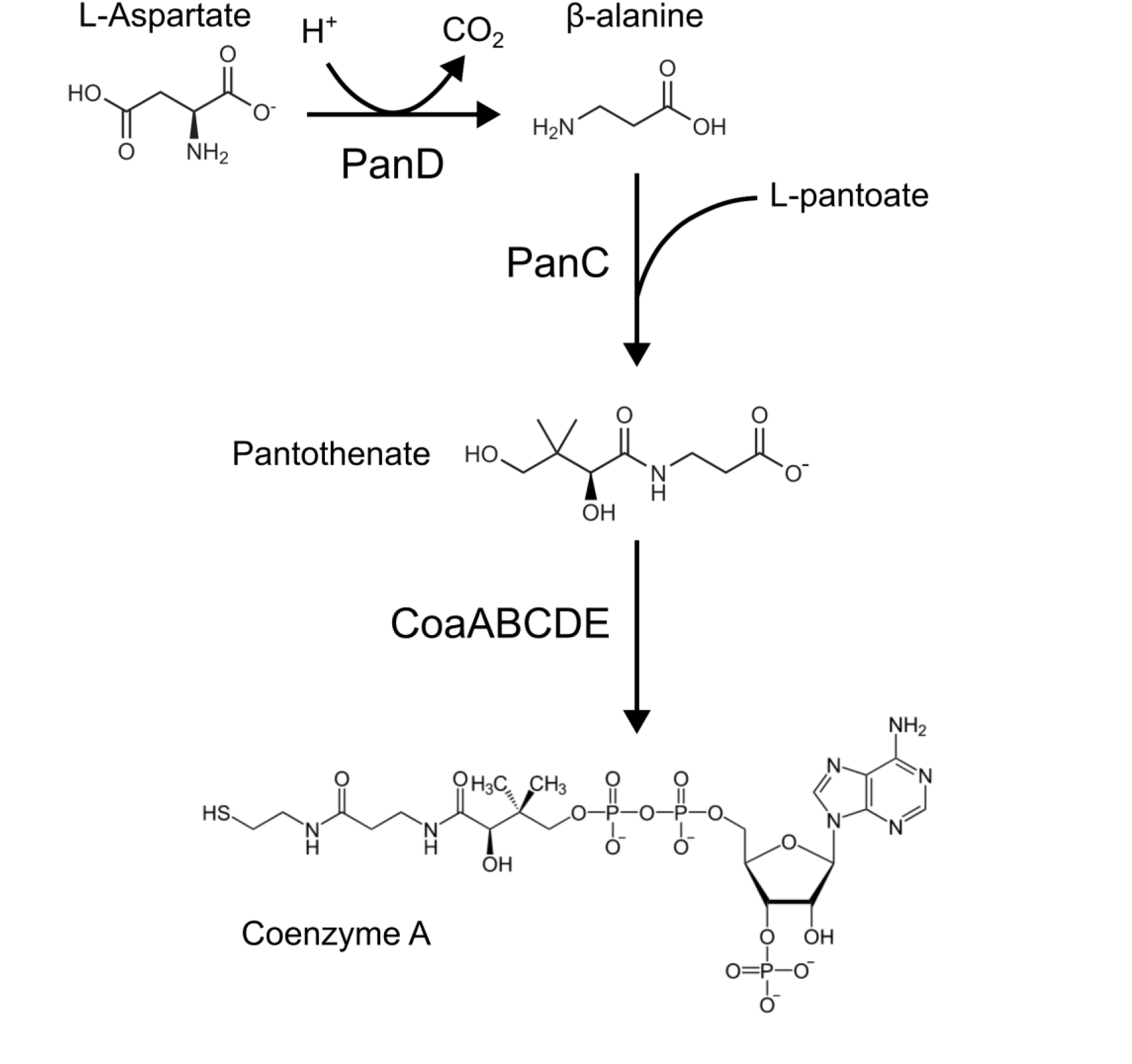
Schematic representation of CoA biosynthesis involving PanCD and CoaABCDE enzymes.

**Figure S2.**
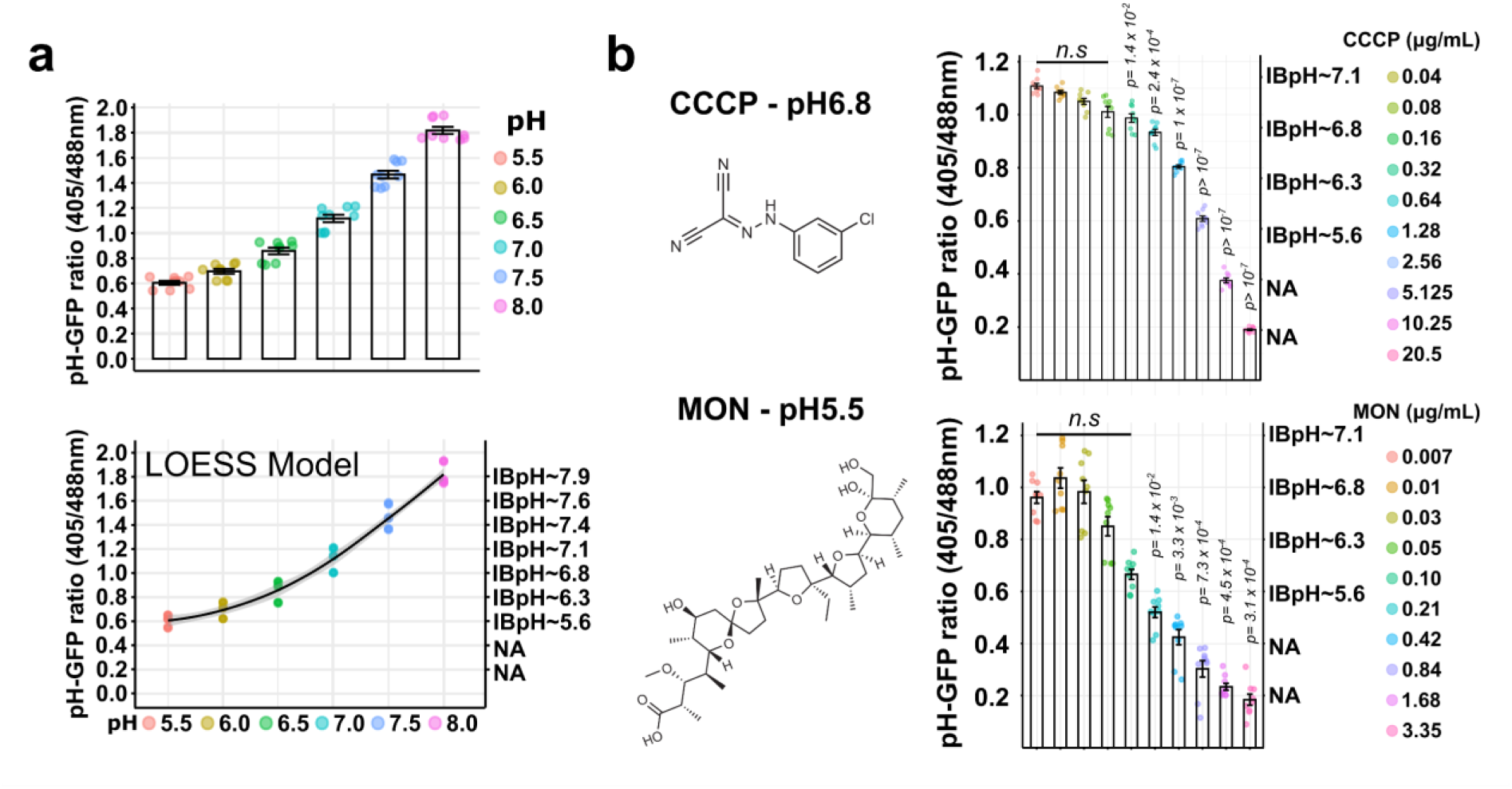
*in vitro* monitoring of *Mtb* IBpH levels and assessment of compound-mediated disruption at both circumneutral and acidic pH. (a) Determination of a calibration curve of *Mtb* H37Ra IBpH homeostasis. *Mtb* pH-GFP cell lysate was resuspended in phosphate citrate buffer with pH values adjusted between pH 5.5 and pH 8. After 48 hours, fluorescence intensity at λ_em_ 535nm recorded after excitation at λ_ex_ 405nm and λ_ex_ 488nm. pH-GFP fluorescence ratios were determined (*top* panel) and used to build a LOESS model that correlates fluorescence intensities with pH values (*bottom* panel). Each measurement is the average of 3 technical replicates performed in 3 biological replicates. **(b)** Quantification of CCCP and Monesin-induced *Mtb* H37Ra IBpH alteration at circum-neutral and acidic pH. *Mtb* pH-GFP ratio were determined in the presence of increasing concentrations of the protonophore CCCP or the pH-dependent ionophore MON after 24 h exposure at pH 6.8 or pH 5.5 respectively. *Mtb* IBpH results displayed in this figure were obtained from 3 biologically independent experiments and are displayed as mean ± SEM. In dose response analysis, statistical significance was assessed by comparing the means of each concentration with the lowest concentration tested using one-way ANOVA followed with Tukey’s multiple comparisons test. All *p*-values were considered significant when *p*-value < 0.05 and their exact values are displayed on the graphs.

**Figure S3.**
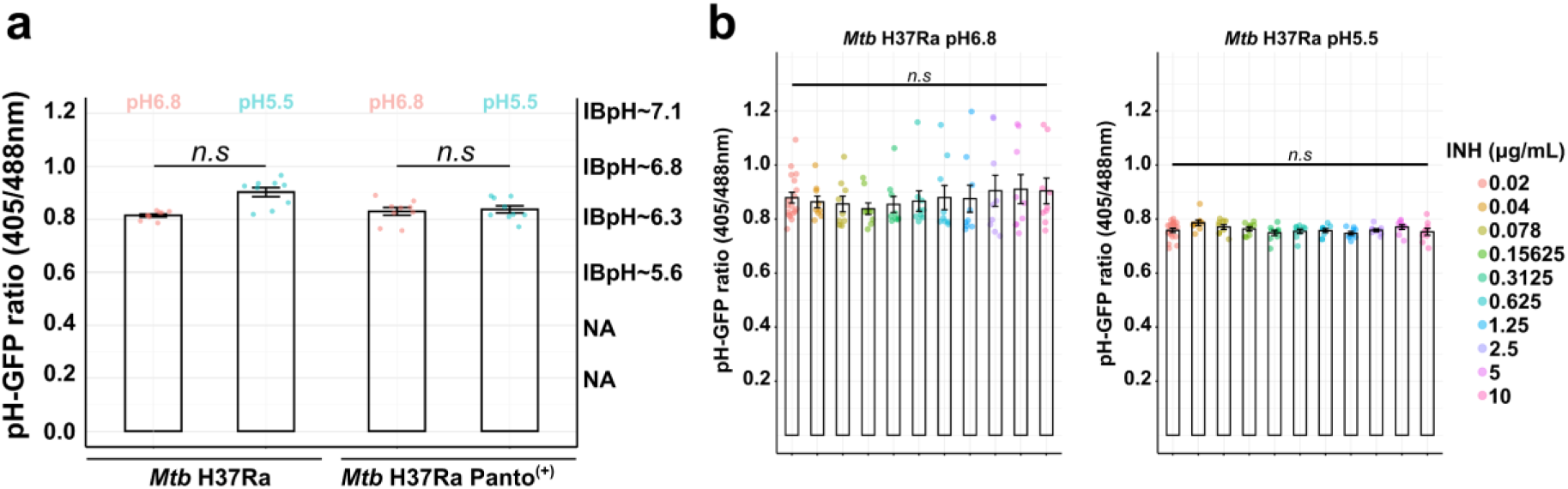
Analysis of *Mtb* IBpH homeostasis maintenance in replicating cells following incubation at circumneutral and acidic pH, in the presence of Panto or in the presence of control drugs. (a) The IBpH of replicating *Mtb* is not significantly affected by external pH or Panto levels. Quantification of *Mtb* H37Ra IBpH alteration was performed at circum-neutral or acidic pH in the absence (*left* panel) or presence (*right* panel) of 25 μg/mL of Panto. Results were obtained after 24 h of incubation in each condition. **(b)** Quantification of INH-induced *Mtb* H37Ra IBpH alteration at circum-neutral (*left* panel) and acidic pH (*right* panel). *Mtb* pH-GFP ratio were determined in the presence of increasing concentrations of control drugs after 24 h of exposure. *Mtb* IBpH results were obtained from 3 biologically independent experiments and are displayed as mean ± SEM. In dose response analysis, statistical significance was assessed by comparing the means of each concentration with the lowest concentration tested using one-way ANOVA followed with Tukey’s multiple comparisons test. All *p*-values were considered significant when *p*-value < 0.05 and their exact values are displayed on the graphs.

**Figure S4.**
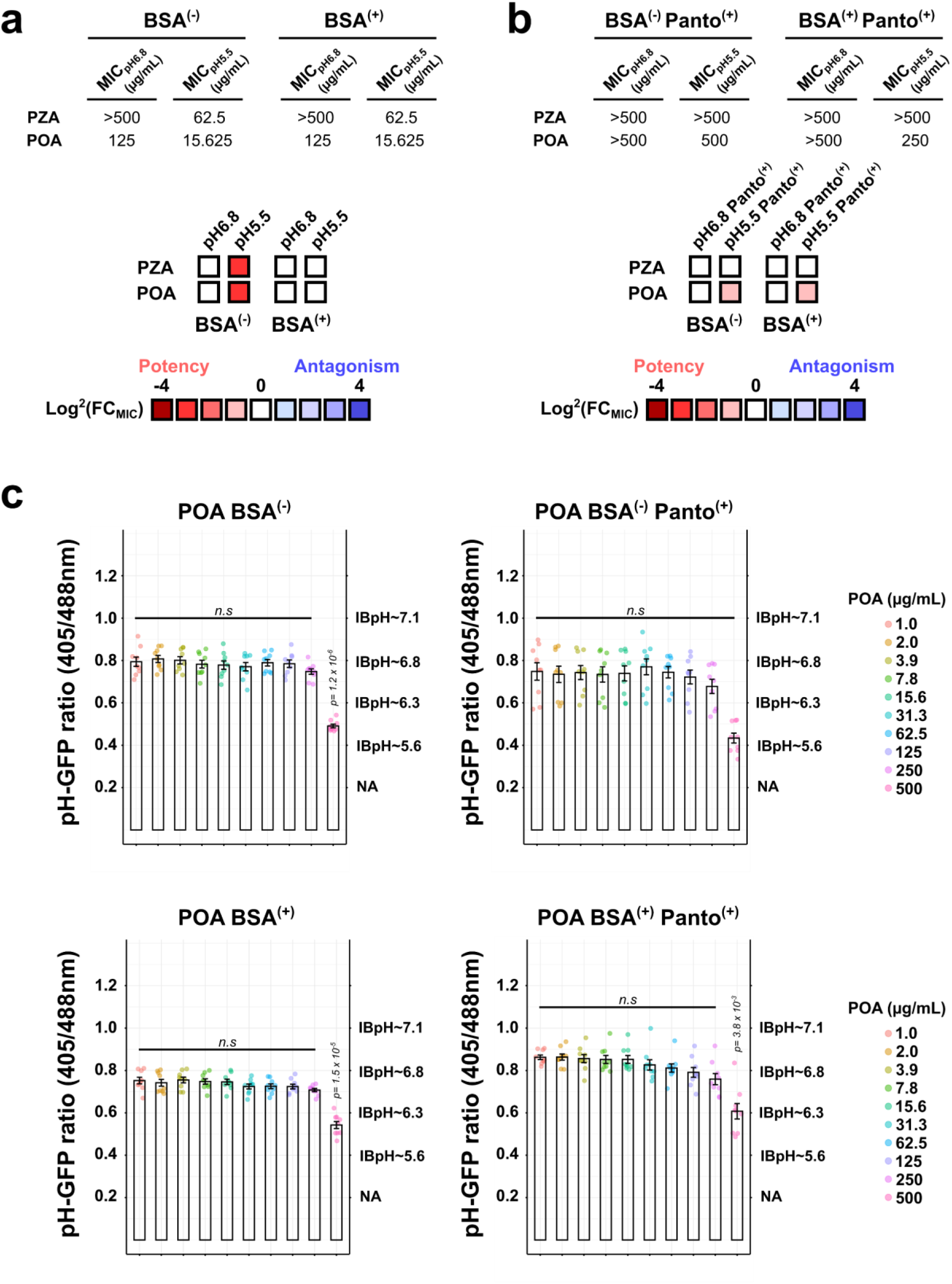
Analysis of BSA’s contribution to POA weak efficacy *in vitro* using antimicrobial susceptibility profiling and monitoring of IBpH homeostasis maintenance in custom made 7H9 media. (a-b) *Mtb* antibiotic susceptibility profiling performed at circum-neutral and acidic pH in the absence (BSA^(-)^) or presence of 5 g/L of BSA (BSA^(+)^) **(a)** without Panto and **(b)** with 25 µg/mL of Panto (Panto^(+)^) . Results are expressed as minimal inhibitory concentration (MIC). Potency and antagonisms are further displayed as color-coded heat map from red (potency) to blue (antagonism). Heat map intensities for each condition represent the fold change (FC) in MIC values expressed as Log^2^(FC_MIC_) by applying the following formula MIC_pH5.5_/MIC_pH6.8_ or MIC_Antagonist_/MIC_CTRL_ at a given pH. All results are representative of three independent technical replicates performed in three biological replicates. **(c)** Quantification of PZA-induced IBpH alteration at acidic pH in *Mtb* H37Ra. *Mtb* pH-GFP ratios were determined in the presence of increasing concentrations of the PZA after 24 h exposure at pH 5.8 in custom Middlebrook 7H9 broth in the presence or absence of 5 g/L of BSA or 25 µg/mL of Panto. All *Mtb* IBpH results displayed in this figure were obtained from 3 biologically independent experiments and are displayed as mean ± SEM. In dose response analysis, statistical significance was assessed by comparing the means of each concentration with the lowest concentration tested using one-way ANOVA followed with Tukey’s multiple comparisons test. All *p*-values were considered significant when *p*-value < 0.05 and their exact values are displayed on the graphs.

**Figure S5.**
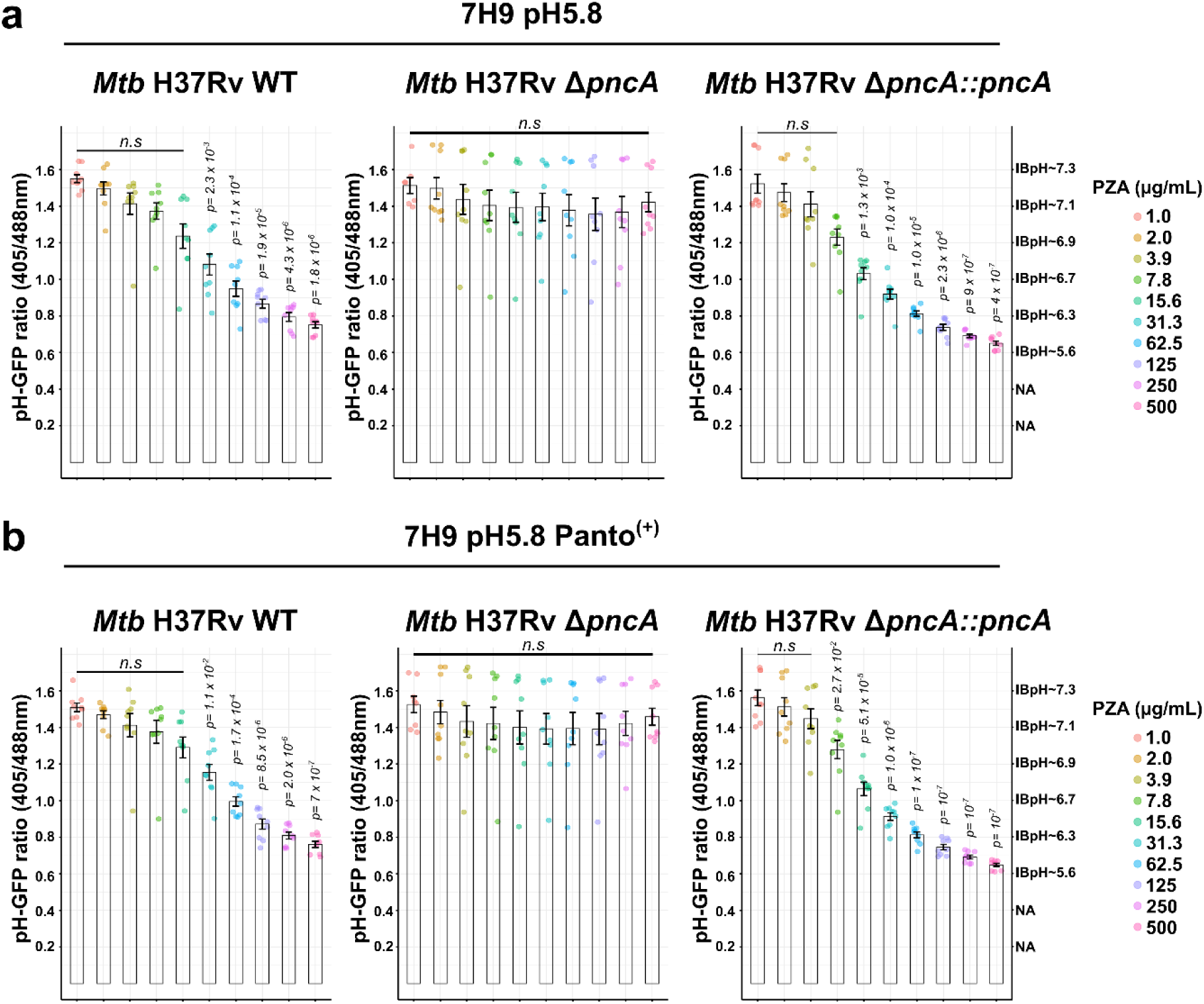
PZA-mediated disruption of IBpH requires a functional *pncA* gene. (a-b) Quantification of PZA-induced IBpH alteration at acidic pH in *Mtb* H37Rv WT, *Mtb* H37Rv Δ*pncA* and *Mtb* H37Rv Δ*pncA::pncA* . *Mtb* pH-GFP ratios were determined in the presence of increasing concentrations of the PZA after 24 h exposure at pH 5.8 in complete Middlebrook 7H9 media **(a)** without Panto and **(b)** with 25 µg/mL of Panto. All *Mtb* IBpH results displayed in this figure were obtained from 3 biologically independent experiments and are displayed as mean ± SEM. In dose response analysis, statistical significance was assessed by comparing the means of each concentration with the lowest concentration tested using one-way ANOVA followed with Tukey’s multiple comparisons test. All *p*-values were considered significant when *p*-value < 0.05 and their exact values are displayed on the graphs.

**Figure S6.**
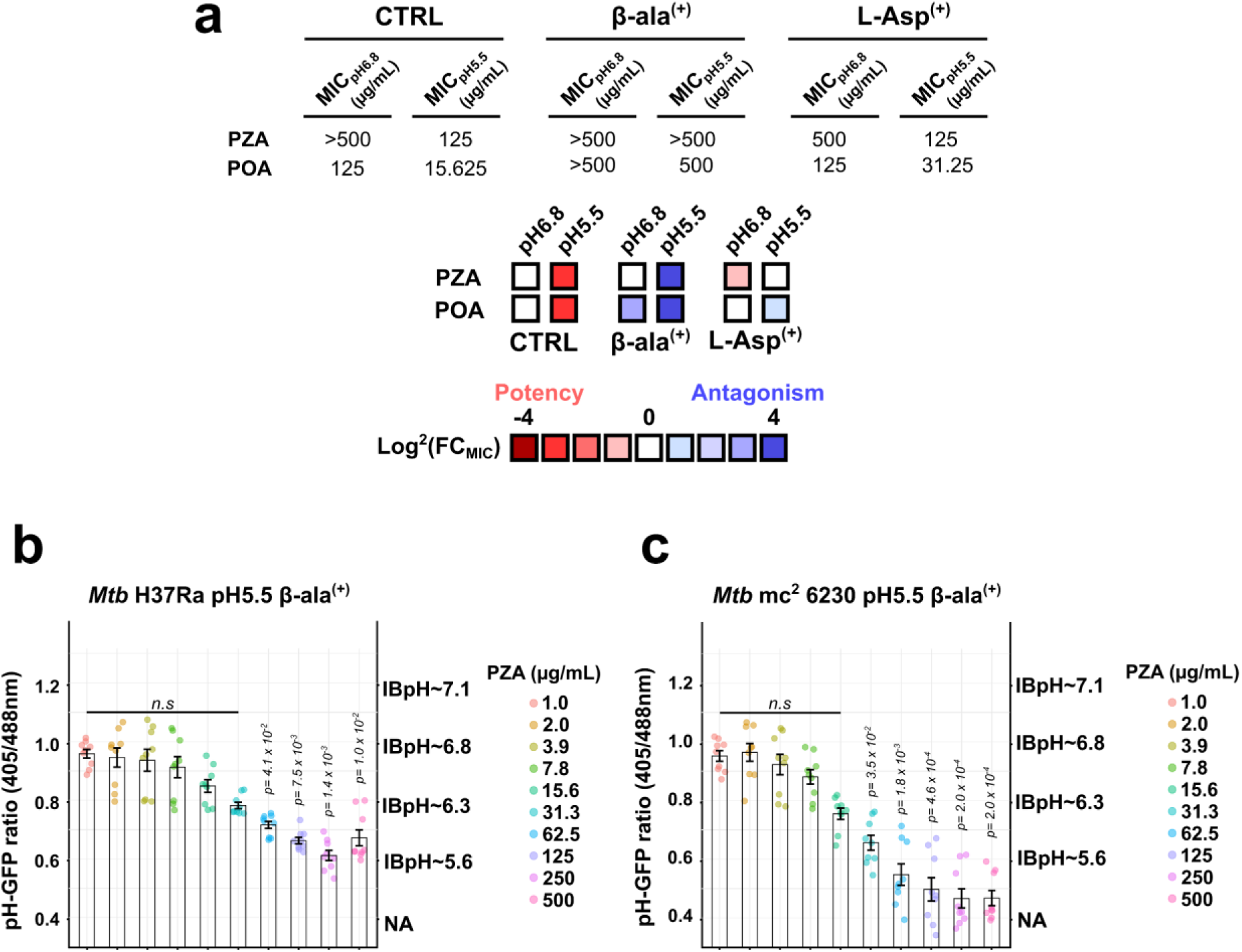
Analysis of β-alanine and L-aspartate antagonistic activity against PZA/POA *in vitro* using antimicrobial susceptibility profiling and monitoring of IBpH homeostasis maintenance. (a) *Mtb* antibiotic susceptibility profiling performed at circum-neutral and acidic pH in the absence (CTRL), presence of β-ala (β-ala^(+)^) or the presence of L-Asp (L-Asp^(+)^). Results are expressed as minimal inhibitory concentration (MIC). Potency and antagonisms are further displayed as color-coded heat map from red (potency) to blue (antagonism). Heat map intensities for each condition represent the fold change (FC) in MIC values expressed as Log^2^(FC_MIC_) by applying the following formula MIC_pH5.5_/MIC_pH6.8_ or MIC_Antagonist_/MIC_CTRL_ at a given pH. All results are representative of three independent technical replicates performed in three biological replicates. **(b)** Quantification of PZA-induced *Mtb* H37Ra IBpH alteration at acidic pH in the presence of 12.5 µg/mL of β-ala (β-ala^(+)^). **(c)** Quantification of PZA-induced *Mtb* mc^2^ 6230 IBpH alteration at acidic pH in the presence of 12.5 µg/mL of β-ala (β-ala^(+)^). *Mtb* pH-GFP ratios were determined in the presence of increasing concentrations of PZA after 24 h of exposure. *Mtb* IBpH results were obtained from 3 biologically independent experiments and are displayed as mean ± SEM. In dose response analysis, statistical significance was assessed by comparing the means of each concentration with the lowest concentration tested using one-way ANOVA followed with Tukey’s multiple comparisons test. All *p*-values were considered significant when *p*-value < 0.05 and their exact values are displayed on the graphs.

**Figure S7.**
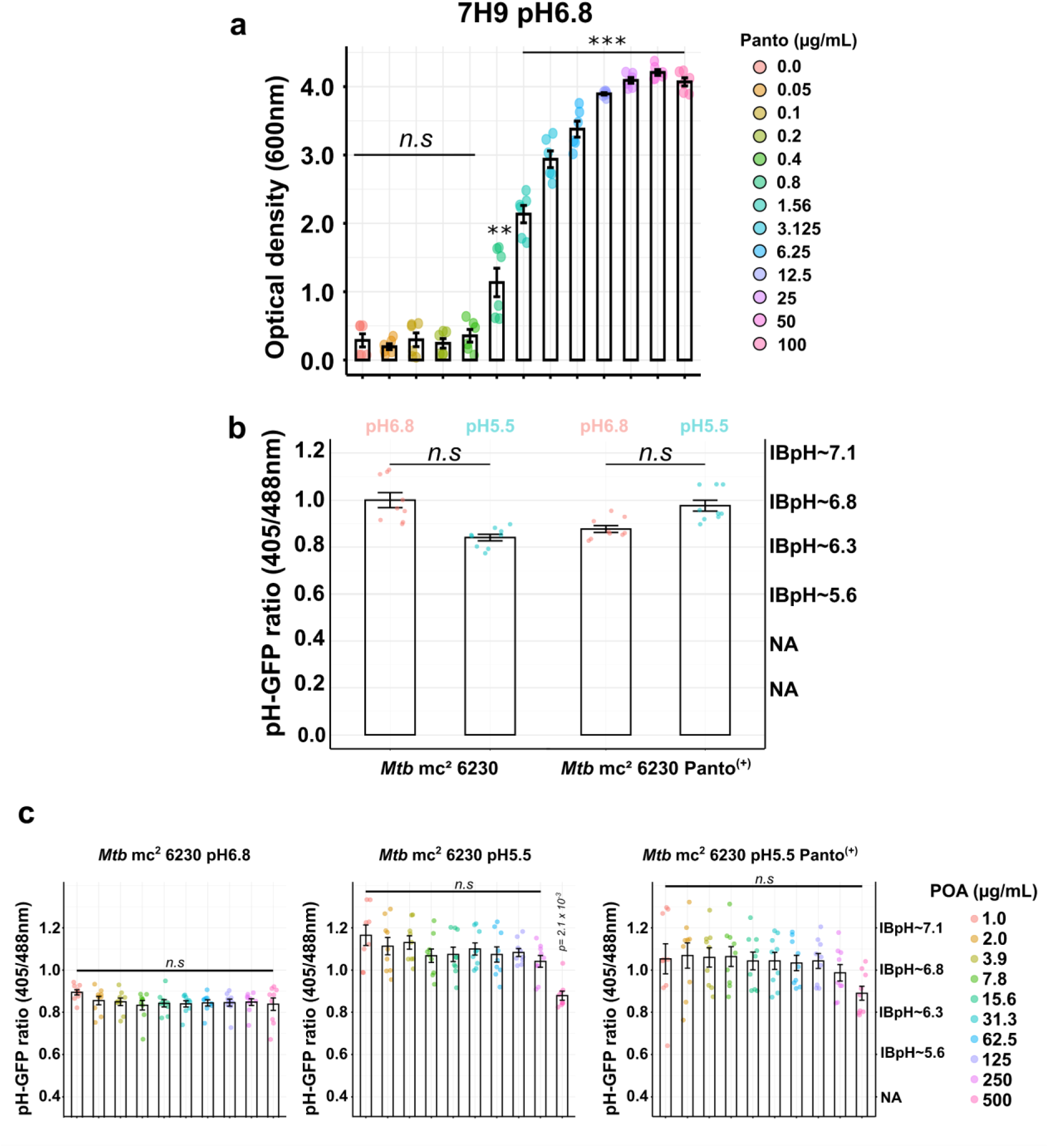
Analysis of *Mtb* mc^2^ 6230 auxotrophy and IBpH homeostasis maintenance in standard 7H9 media following incubation at circumneutral and acidic pH. (a) Panto auxotrophy experiments confirming the non-growing phenotype of *Mtb* mc^2^ 6230 in the absence of exogenous Panto. A dose response analysis of Panto supplementation was performed confirming that concentrations greater than 0.8 µg/mL support replication, with optimal growth observed when using 25 µg/mL and beyond. **(b)** The IBpH of *Mtb* mc^2^ 6230 is not significantly affected by external pH or Panto levels. Quantification of *Mtb* mc^2^ 6230 IBpH alteration were performed at circum-neutral or acidic pH in the absence (*left* panel) or presence (*right* panel) of 25 µg/mL of Panto. Results were obtained after 24 h of incubation in each condition. **(c)** Quantification of POA-induced *Mtb* H37Ra IBpH alteration at circum-neutral (*left* panel), acidic pH (*middle* panel) and acidic pH in the presence of 25 µg/mL of Panto (*right* panel). *Mtb* pH-GFP ratios were determined in the presence of increasing concentrations of POA after 24 h of exposure. *Mtb* IBpH results were obtained from 3 biologically independent experiments and are displayed as mean ± SEM. In dose response analysis, statistical significance was assessed by comparing the means of each concentration with the lowest concentration tested using one-way ANOVA followed with Tukey’s multiple comparisons test. All *p*-values were considered significant when *p*-value < 0.05 and their exact values are displayed on the graphs.

**Figure S8.**
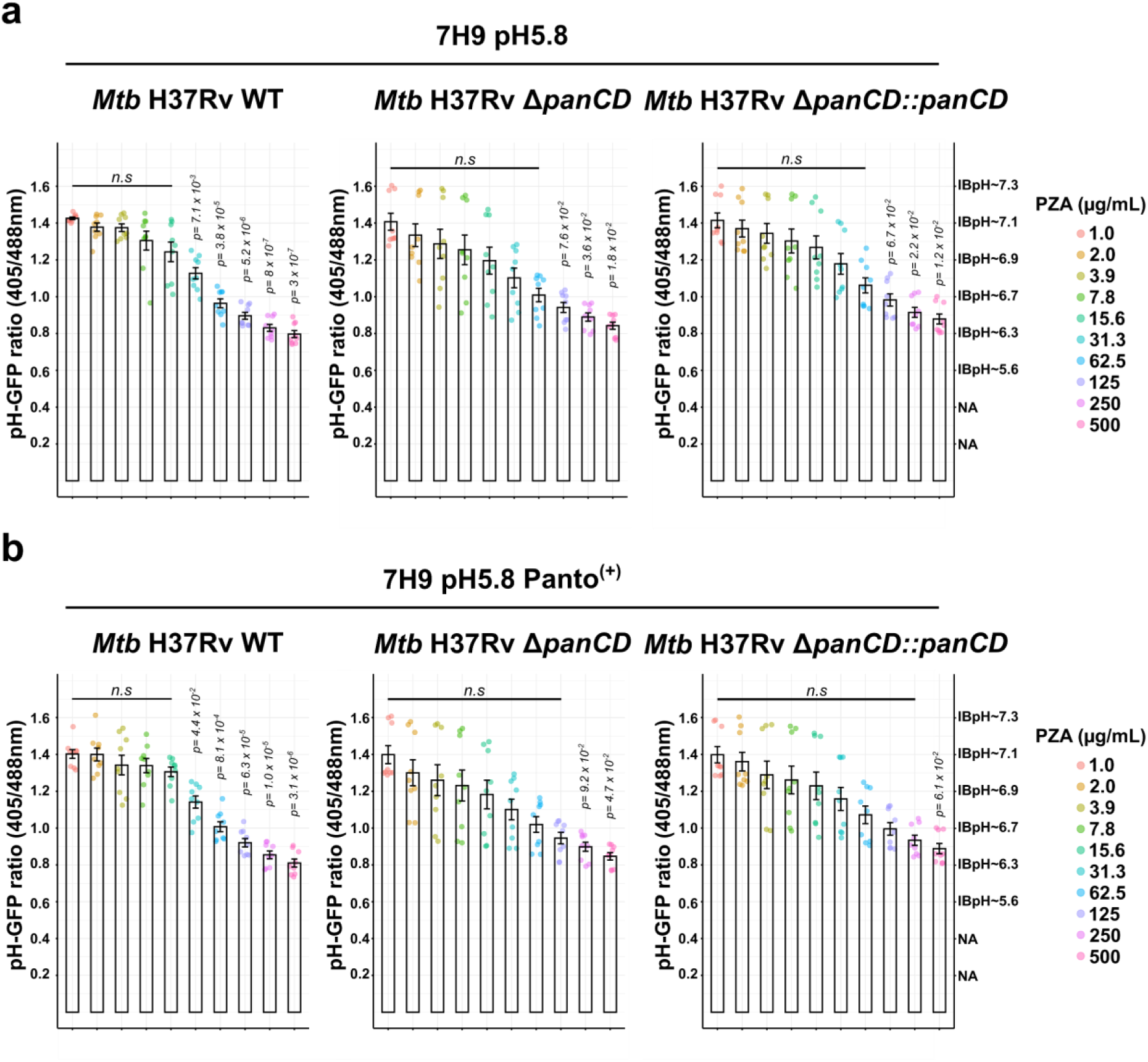
PZA-mediated disruption of IBpH is not impacted by the lack of *panCD* locus nor supplementation of Panto while assessed with H37Rv-derived isogenic strains. (a-b) Quantification of PZA-induced IBpH alteration at acidic pH in *Mtb* H37Rv WT, *Mtb* H37Rv Δ*panCD* and *Mtb* H37Rv Δ*panCD::panCD* . *Mtb* pH-GFP ratios were determined in the presence of increasing concentrations of the PZA after 24 h exposure at pH 5.8 in complete Middlebrook 7H9 media **(a)** without Panto and **(b)** with 25 µg/mL of Panto. All *Mtb* IBpH results displayed in this figure were obtained from 3 biologically independent experiments and are displayed as mean ± SEM. In dose response analysis, statistical significance was assessed by comparing the means of each concentration with the lowest concentration tested using one-way ANOVA followed with Tukey’s multiple comparisons test. All *p*-values were considered significant when *p*-value < 0.05 and their exact values are displayed on the graphs.

**Figure S9.**
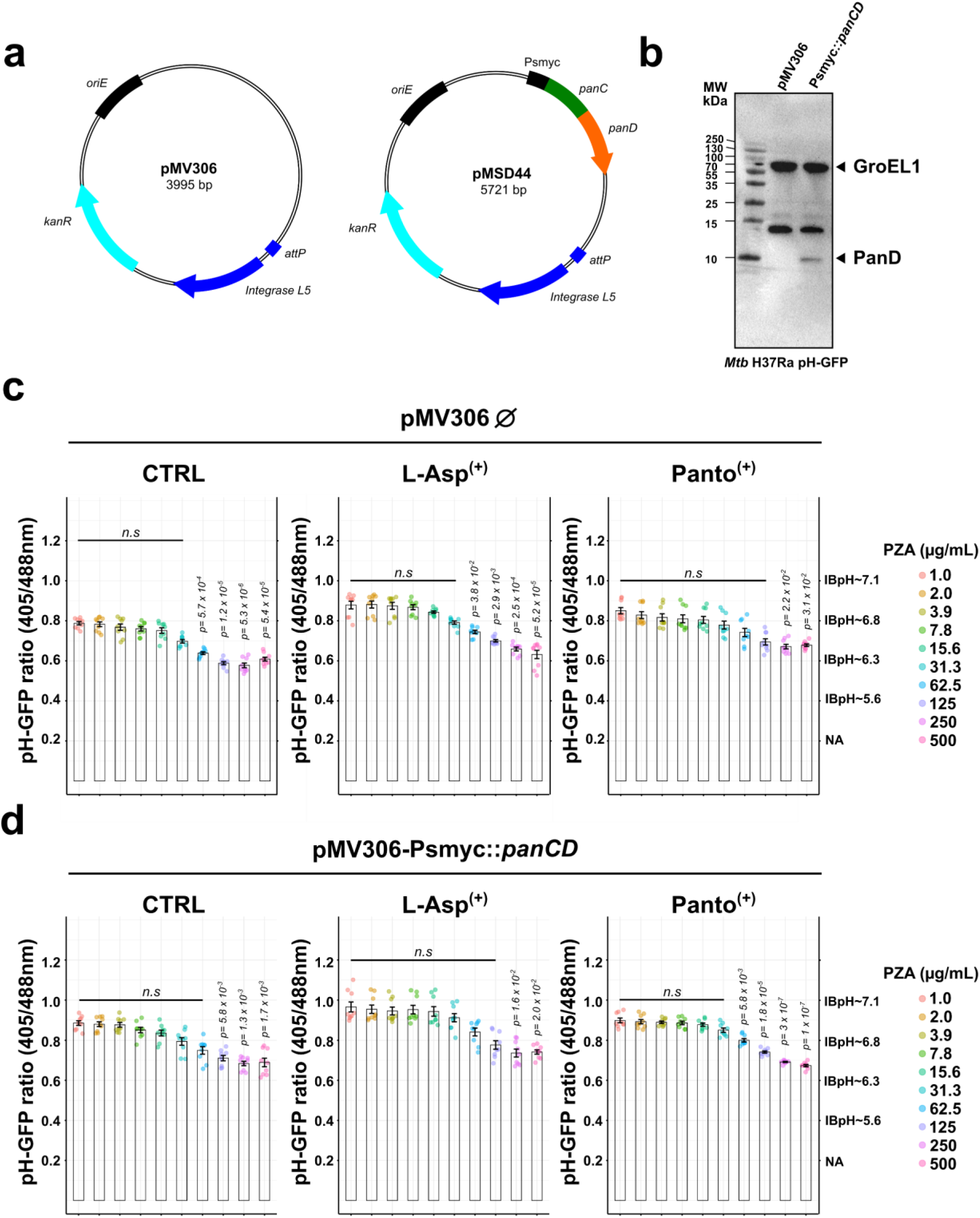
*panCD* overexpression does not prevent from PZA-induced IBpH acidification. **(a)** Schematic representation of the plasmid construct used to overexpress *panCD*. Empty pMV306 was used as control while pMV306-Psmyc::*panCD* also referred to pMSD44 was used as an integrative vector to overexpress the *panCD* locus in *cis*. **(b)** Immunoblot analysis of 6xHis tagged protein levels in *Mtb* H37Ra pH-GFP, highlighting PanD production. Normalized samples were loaded onto 15% SDS/PAGE and immunoblotted using HisProbe® reagent. Black arrows represent the 5-His containing protein GroEL1 and PanD respectively. **(c)** Quantification of PZA-induced IBpH alteration at acidic pH in *Mtb* H37Ra pH-GFP pMV306 or **(d)** in *Mtb* H37Ra pH-GFP pMV306-Psmyc::*panCD*. *Mtb* pH-GFP ratios were determined in the presence of increasing concentrations of PZA after 24 h exposure at pH 5.8 in complete Middlebrook 7H9 media (CTRL - *left* panel); or supplemented with L-Asp (L-Asp^(+)^ - *middle* panel) or supplemented with Panto (Panto^(+)^ - *right* panel). All *Mtb* IBpH results displayed in this figure were obtained from 3 biologically independent experiments and are displayed as mean ± SEM. In dose response analysis, statistical significance was assessed by comparing the means of each concentration with the lowest concentration tested using one-way ANOVA followed with Tukey’s multiple comparisons test. All *p*-values were considered significant when *p*-value < 0.05 and their exact values are displayed on the graphs.

**Figure S10.**
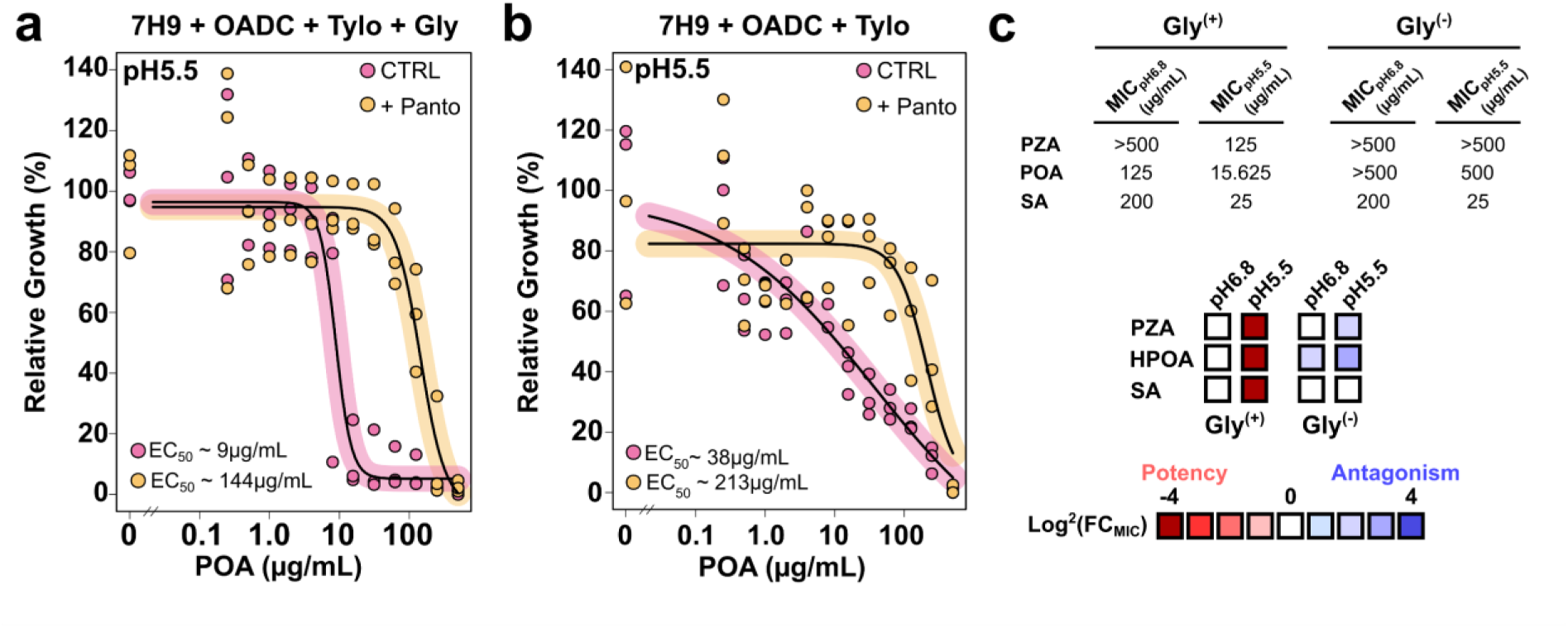
PZA/POA efficacy and Panto-mediated antagonism are impacted by carbon sources. **(a-b)** Comparative antibiotic susceptibility of *Mtb* H37Ra towards POA performed at mildly acidic pH in 7H9+OADC+Tyloxapol culture media that contains (**a** – *left* panel) or not 0.2% glycerol (**b** – *right* panel). Experiments were performed in the absence (pink) or presence of Panto (yellow) antagonist. Results are expressed as relative growth % where the no drug condition represents 100%. Dose-response curves displayed were obtained following a 4-parameter nonlinear logistic regression and EC_50_ were determined accordingly. **(c)** Summary of glycerol effect on PZA/POA efficacy and antagonisms. Results are expressed as minimal inhibitory concentration (MIC) and further displayed as color-coded heatmap from red (potency) to blue (antagonism). Heatmap intensities for each compound represent the fold change (FC) in MIC values expressed as Log^2^(FC_MIC_) by applying the following formula MIC_pH5.5_/MIC_pH6.8_ or MIC_Antagonist_/MIC_CTRL_. Results are representative of three independent replicates performed in biological replicates.

**Figure S11.**
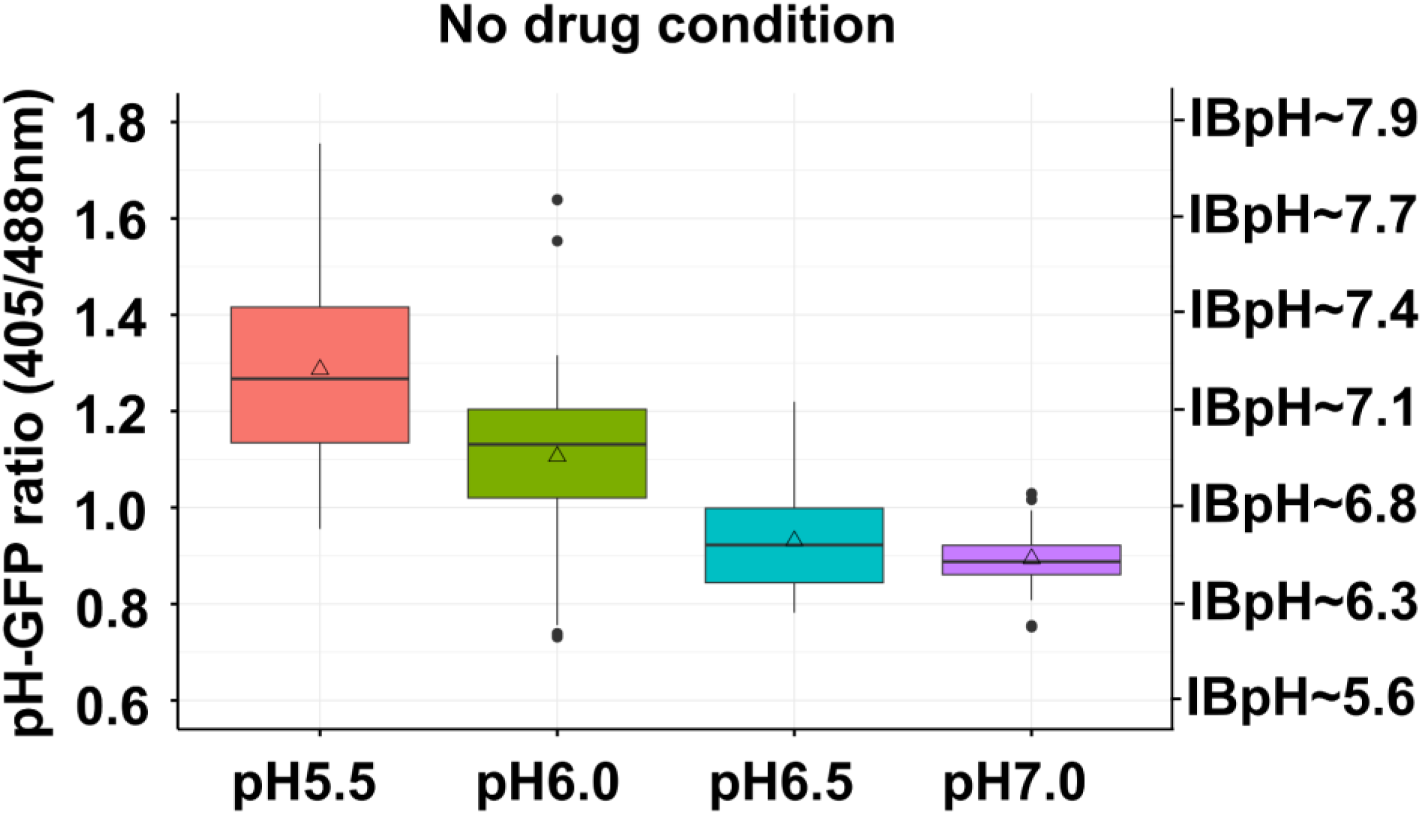
Analysis of *Mtb* IBpH homeostasis maintenance in non-replicating starved cells following incubation in phosphate citrate buffer at different pH. pH-dependent analysis of *Mtb* H37Ra IBpH homeostasis. pH-GFP ratios were determined after a 48 h incubation period in phosphate citrate buffer adjusted at pH 7 (purple), pH 6.5 (blue), pH 6 (green) and pH 5.5 (red).

**Figure S12.**
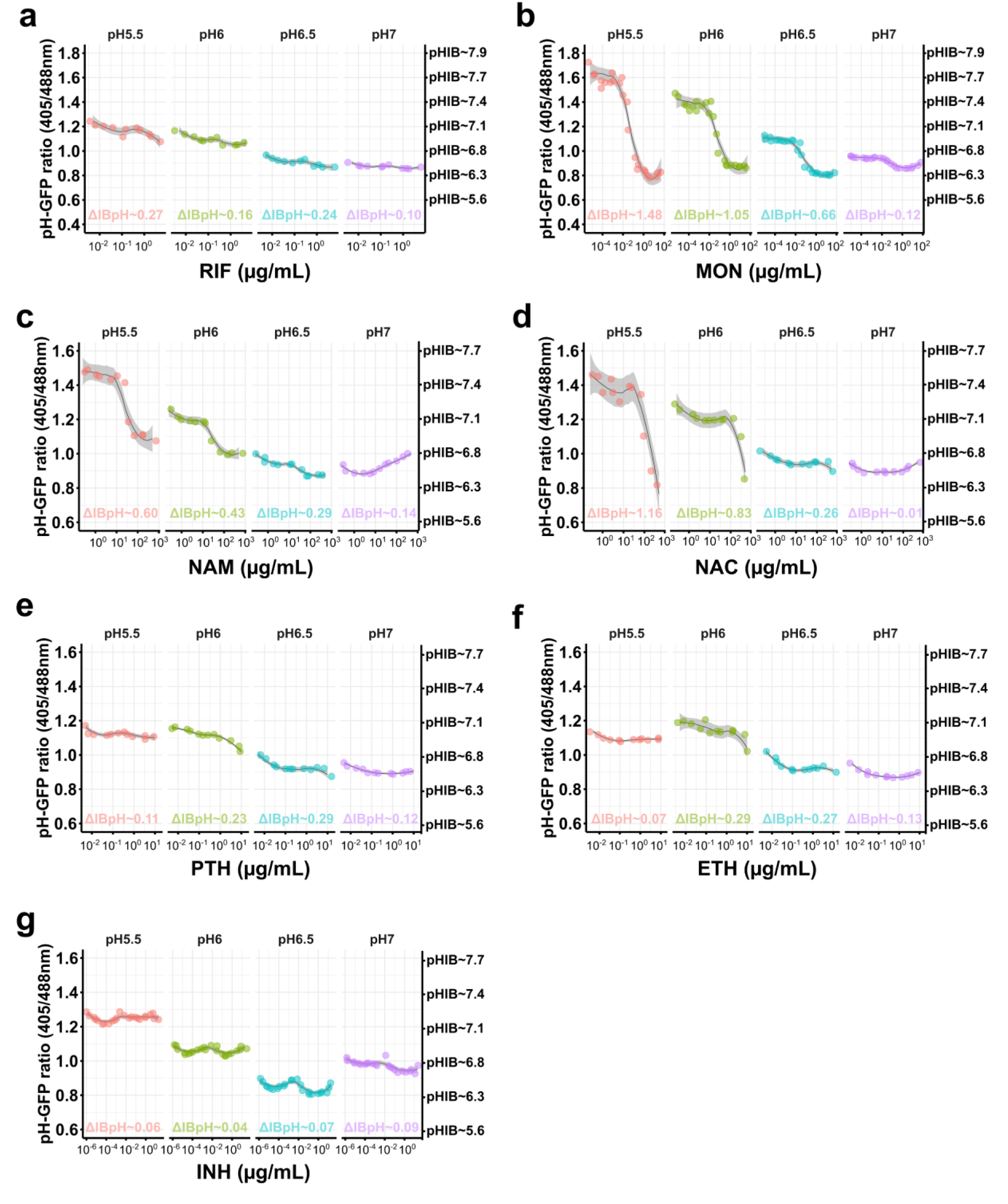
Analysis of *Mtb* IBpH homeostasis in non-replicating cells upon treatment with a large panel of drugs. **(a-g)** pH-dependent effect of RIF (**a**), MON (**b**), NAM (**c**), NAC (**d**), PTH (**e**), ETH (**f**) and INH (**g**) on *Mtb* H37Ra IBpH homeostasis at distinct pH. Dose-response analyses were performed in phosphate citrate buffer adjusted at pH 7 (purple), pH 6.5 (blue), pH 6 (green) and pH 5.5 (red). *Mtb* were exposed to increasing concentrations of RIF or MON for 48 h before performing pH-GFP ratio recording and determination.

**Figure S13.**
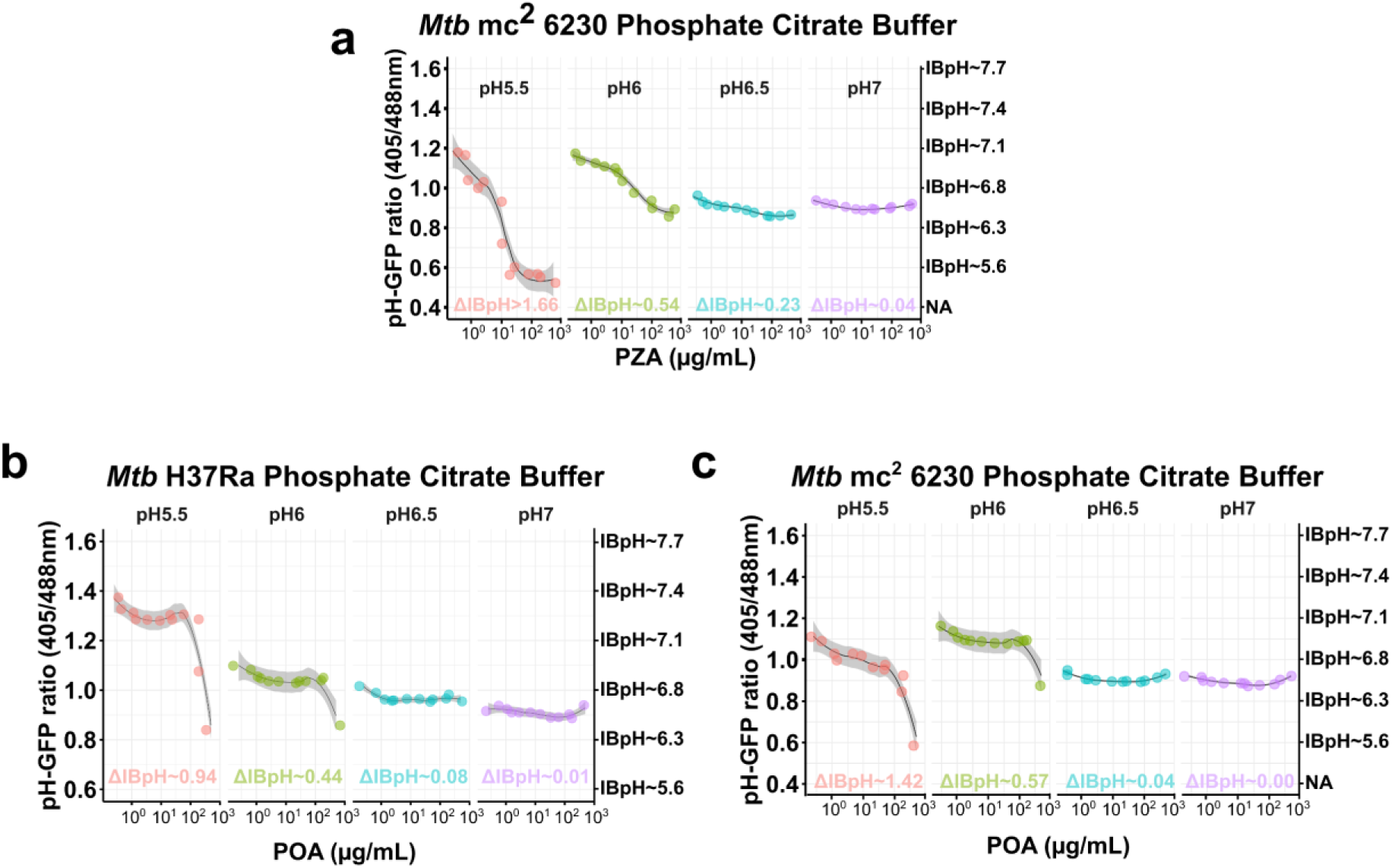
Analysis of *Mtb* IBpH homeostasis in non-replicating cells upon treatment with PZA/POA. **(a)** pH-dependent PZA-mediated IBpH cytosolic acidification of *Mtb* mc^2^ 6230. Dose-response analyses were performed in phosphate citrate buffer adjusted at pH 7 (purple), pH 6.5 (blue), pH 6 (green) and pH 5.5 (red). *Mtb* were exposed to increasing concentrations of PZA from 0.25 to 500 µg/mL for 48 h before pH-GFP ratio recording. **(b-c)** pH-dependent POA-mediated IBpH cytosolic acidification of *Mtb* H37Ra and *Mtb* mc^2^ 6230. Dose-response analyses were performed in phosphate citrate buffer adjusted at pH 7 (purple), pH 6.5 (blue), pH 6 (green) and pH 5.5 (red). *Mtb* were exposed to increasing concentrations of POA from 0.25 to 500 µg/mL for 48 h before pH-GFP ratio recording. Using pH-GFP ratios, distinct models were built using the LOESS function and ΔIBpH were calculated by comparing the IBpH recorded at the highest POA concentration tested with the one at the lowest POA concentration included in the dose response for each tested pH.

**Figure S14.**
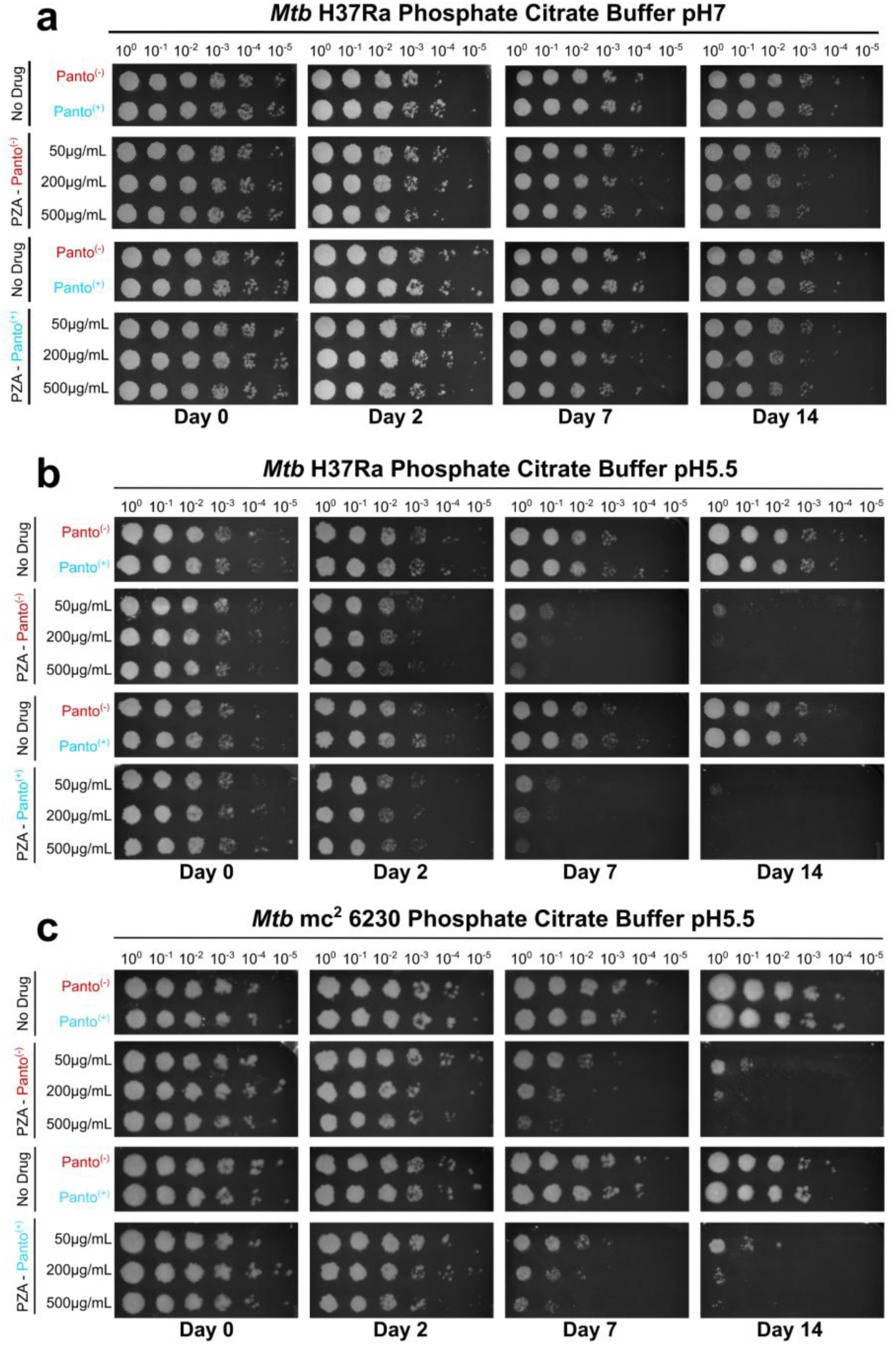
Acidic pH governs PZA’s ability kill non-replicating *Mtb* independently of the *panCD* locus and pantothenate levels. **(a-b-c)** pH-dependent bactericidal activity of PZA on non-replicating *Mtb* H37Ra and *Mtb* mc^2^ 6230 pH-GFP cells. Survival in phosphate citrate buffer was performed by pulsing 1 x 10^7^ *Mtb* CFU/mL with increasing concentration of PZA including 0 μg/mL, 50 μg/mL, 200 μg/mL and 500 μg/mL in the presence or absence of 25 µg/mL Panto. At day 0, 2, 7 and 14, *Mtb* cells were recovered, serially diluted and spotted onto 7H11 agar media in the presence 25 µg/mL Panto and 50 µg/mL hygromycin B. Experiments were performed with *Mtb* H37Ra at pH 7 (**a** – *top* panel) and at pH 5.5 (**b** – *middle* panel), and with *Mtb* mc^2^ 6230 at pH 5.5 (**c** – *bottom* panel). Results are representative of two distinct experiments performed on biological independent occasion.

**Figure S15.**
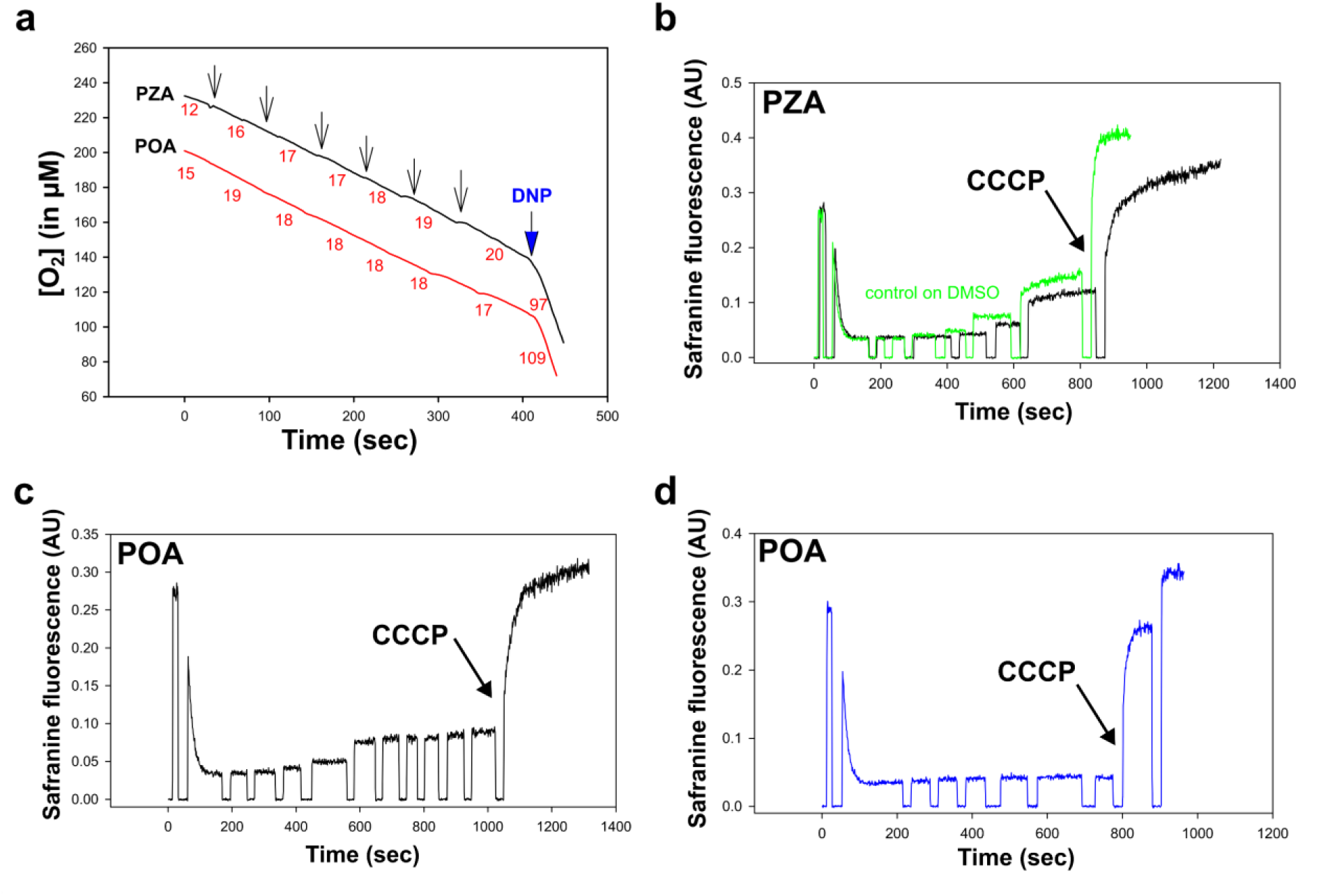
PZA/POA has no effect on mitochondrial respiration and membrane potential. **(a)** Effect of PZA and POA on respiration of rat liver mitochondria (RLM). The substrate was succinate supplemented with 2 μM rotenone. Incubation medium contained 250 mM sucrose, 20 mM MOPS, 1 mM KH_2_PO_4_, 2 mM MgCl_2_, 1 mM EGTA, pH 7.2, and mitochondria (0.5 mg protein/mL). Numbers at different parts of each record are respiration rates in relative units. At the ned of the kinetic, 40 μM DNP was added as positive control. **(b-c-d)** Effect of PZA (**b**) and POA (**c-d**) on mitochondrial membrane potential estimated with the fluorescence of potential-sensitive dye safranine O (15 µM) in isolated rat liver mitochondria (RLM). Y-axis shows fluorescence of safranine at 580 nm (λ_ext_ 520 nm). Increasing volume 2 µL, 4 µL, 10 µL, 20 µL and 40 µL of PZA, POA or DMSO were added to sample and changes in safranine O fluorescence were recorded. The substrate was succinate supplemented with 2 μM rotenone. 200 nM CCCP was added as positive control at the end of the kinetics.

## References

1. (WHO) WHO. Global tuberculosis report 2025. .) (2025).

2. Dartois VA, Rubin EJ. Anti-tuberculosis treatment strategies and drug development: challenges and priorities. Nat Rev Microbiol 20, 685–701 (2022).

3. Malone L, Schurr A, Lindh H, Mc KD, Kiser JS, Williams JH. The effect of pyrazinamide (aldinamide) on experimental tuberculosis in mice. Am Rev Tuberc 65, 511–518 (1952).

4. Solotorovsky M, Gregory FJ, Ironson EJ, Bugie EJ, O’Neill RC, Pfister R, 3rd. Pyrazinoic acid amide; an agent active against experimental murine tuberculosis. Proc Soc Exp Biol Med 79, 563–565 (1952).

5. Tarshis MS, Weed WA, Jr. Lack of significant in vitro sensitivity of Mycobacterium tuberculosis to pyrazinamide on three different solid media. Am Rev Tuberc 67, 391–395 (1953).

6. Mitchison DA. The action of antituberculosis drugs in short-course chemotherapy. Tubercle 66, 219–225 (1985).

7. Zhang Y, Shi W, Zhang W, Mitchison D. Mechanisms of Pyrazinamide Action and Resistance. Microbiol Spectr 2, 1–12 (2013).

8. Guglielmetti L, et al. Oral Regimens for Rifampin-Resistant, Fluoroquinolone-Susceptible Tuberculosis. N Engl J Med 392, 468–482 (2025).

9. Kushner S, et al. Experimental chemotherapy of tuberculosis. II. The synthesis of pyrazinamides and related compounds1. 74, 3617–3621 (1952).

10. Yeager RL, Munroe WG, Dessau FI. Pyrazinamide (aldinamide*) in the treatment of pulmonary tuberculosis. Trans Annu Meet Natl Tuberc Assoc 48, 178–201 (1952).

11. McCune RM, Jr., Tompsett R. Fate of Mycobacterium tuberculosis in mouse tissues as determined by the microbial enumeration technique. I. The persistence of drug-susceptible tubercle bacilli in the tissues despite prolonged antimicrobial therapy. J Exp Med 104, 737–762 (1956).

12. Fox W, Ellard GA, Mitchison DA. Studies on the treatment of tuberculosis undertaken by the British Medical Research Council tuberculosis units, 1946-1986, with relevant subsequent publications. Int J Tuberc Lung Dis 3, S231–279 (1999).

13. Gazzaniga F, Stebbins R, Chang SZ, McPeek MA, Brenner C. Microbial NAD metabolism: lessons from comparative genomics. Microbiol Mol Biol Rev 73, 529–541, Table of Contents (2009).

14. Boshoff HI, et al. Biosynthesis and recycling of nicotinamide cofactors in mycobacterium tuberculosis. An essential role for NAD in nonreplicating bacilli. J Biol Chem 283, 19329–19341 (2008).

15. Zignol M, et al. Population-based resistance of Mycobacterium tuberculosis isolates to pyrazinamide and fluoroquinolones: results from a multicountry surveillance project. Lancet Infect Dis 16, 1185–1192 (2016).

16. Scorpio A, Zhang Y. Mutations in pncA, a gene encoding pyrazinamidase/nicotinamidase, cause resistance to the antituberculous drug pyrazinamide in tubercle bacillus. Nat Med 2, 662–667 (1996).

17. Consortium CR, et al. Prediction of Susceptibility to First-Line Tuberculosis Drugs by DNA Sequencing. N Engl J Med 379, 1403–1415 (2018).

18. The CC. A data compendium associating the genomes of 12,289 Mycobacterium tuberculosis isolates with quantitative resistance phenotypes to 13 antibiotics. PLoS Biol 20, e3001721 (2022).

19. Zimhony O, Cox JS, Welch JT, Vilcheze C, Jacobs WR, Jr. Pyrazinamide inhibits the eukaryotic-like fatty acid synthetase I (FASI) of Mycobacterium tuberculosis. Nat Med 6, 1043–1047 (2000).

20. Shi W, et al. Pyrazinamide inhibits trans-translation in Mycobacterium tuberculosis. Science 333, 1630–1632 (2011).

21. Boshoff HI, Mizrahi V, Barry CE, 3rd. Effects of pyrazinamide on fatty acid synthesis by whole mycobacterial cells and purified fatty acid synthase I. J Bacteriol 184, 2167–2172 (2002).

22. Dillon NA, Peterson ND, Feaga HA, Keiler KC, Baughn AD. Anti-tubercular Activity of Pyrazinamide is Independent of trans-Translation and RpsA. Sci Rep 7, 6135 (2017).

23. Salfinger M, Heifets LB. Determination of pyrazinamide MICs for Mycobacterium tuberculosis at different pHs by the radiometric method. Antimicrob Agents Chemother 32, 1002–1004 (1988).

24. Zhang Y, Scorpio A, Nikaido H, Sun Z. Role of acid pH and deficient efflux of pyrazinoic acid in unique susceptibility of Mycobacterium tuberculosis to pyrazinamide. J Bacteriol 181, 2044–2049 (1999).

25. Zhang Y, Permar S, Sun Z. Conditions that may affect the results of susceptibility testing of Mycobacterium tuberculosis to pyrazinamide. J Med Microbiol 51, 42–49 (2002).

26. Zhang Y, Wade MM, Scorpio A, Zhang H, Sun Z. Mode of action of pyrazinamide: disruption of Mycobacterium tuberculosis membrane transport and energetics by pyrazinoic acid. J Antimicrob Chemother 52, 790–795 (2003).

27. Zhang Y, Mitchison D. The curious characteristics of pyrazinamide: a review. Int J Tuberc Lung Dis 7, 6–21 (2003).

28. Fontes FL, Peters BJ, Crans DC, Crick DC. The Acid-Base Equilibrium of Pyrazinoic Acid Drives the pH Dependence of Pyrazinamide-Induced Mycobacterium tuberculosis Growth Inhibition. ACS Infect Dis 6, 3004–3014 (2020).

29. Darby CM, et al. Whole cell screen for inhibitors of pH homeostasis in Mycobacterium tuberculosis. PLoS One 8, e68942 (2013).

30. Santucci P, et al. Visualizing Pyrazinamide Action by Live Single-Cell Imaging of Phagosome Acidification and Mycobacterium tuberculosis pH Homeostasis. mBio 13, e0011722 (2022).

31. Zhang S, Chen J, Shi W, Liu W, Zhang W, Zhang Y. Mutations in panD encoding aspartate decarboxylase are associated with pyrazinamide resistance in Mycobacterium tuberculosis. Emerg Microbes Infect 2, e34 (2013).

32. Shi W, et al. Aspartate decarboxylase (PanD) as a new target of pyrazinamide in Mycobacterium tuberculosis. Emerg Microbes Infect 3, e58 (2014).

33. Gopal P, et al. Pyrazinamide Resistance Is Caused by Two Distinct Mechanisms: Prevention of Coenzyme A Depletion and Loss of Virulence Factor Synthesis. ACS Infect Dis 2, 616–626 (2016).

34. den Hertog AL, Menting S, Pfeltz R, Warns M, Siddiqi SH, Anthony RM. Pyrazinamide Is Active against Mycobacterium tuberculosis Cultures at Neutral pH and Low Temperature. Antimicrob Agents Chemother 60, 4956–4960 (2016).

35. Peterson ND, Rosen BC, Dillon NA, Baughn AD. Uncoupling Environmental pH and Intrabacterial Acidification from Pyrazinamide Susceptibility in Mycobacterium tuberculosis. Antimicrob Agents Chemother 59, 7320–7326 (2015).

36. Sun Q, Li X, Perez LM, Shi W, Zhang Y, Sacchettini JC. The molecular basis of pyrazinamide activity on Mycobacterium tuberculosis PanD. Nat Commun 11, 339 (2020).

37. Gopal P, et al. Pyrazinamide triggers degradation of its target aspartate decarboxylase. Nat Commun 11, 1661 (2020).

38. Gopal P, et al. Pyrazinoic Acid Inhibits Mycobacterial Coenzyme A Biosynthesis by Binding to Aspartate Decarboxylase PanD. ACS Infect Dis 3, 807–819 (2017).

39. Gopal P, Gruber G, Dartois V, Dick T. Pharmacological and Molecular Mechanisms Behind the Sterilizing Activity of Pyrazinamide. Trends Pharmacol Sci 40, 930–940 (2019).

40. Lamont EA, Dillon NA, Baughn AD. The Bewildering Antitubercular Action of Pyrazinamide. Microbiol Mol Biol Rev 84, (2020).

41. Lamont EA, Baughn AD. Impact of the host environment on the antitubercular action of pyrazinamide. EBioMedicine 49, 374–380 (2019).

42. Anthony RM, den Hertog AL, van Soolingen D. ’Happy the man, who, studying nature’s laws, Thro’ known effects can trace the secret cause.’ Do we have enough pieces to solve the pyrazinamide puzzle? J Antimicrob Chemother 73, 1750–1754 (2018).

43. Laudouze J, et al. Antitubercular potential and pH-driven mode of action of salicylic acid derivatives. FEBS Open Bio 15, 383–398 (2025).

44. Zhang Y, Zhang H, Sun Z. Susceptibility of Mycobacterium tuberculosis to weak acids. J Antimicrob Chemother 52, 56–60 (2003).

45. Dillon NA, Peterson ND, Rosen BC, Baughn AD. Pantothenate and pantetheine antagonize the antitubercular activity of pyrazinamide. Antimicrob Agents Chemother 58, 7258–7263 (2014).

46. Vandal OH, Pierini LM, Schnappinger D, Nathan CF, Ehrt S. A membrane protein preserves intrabacterial pH in intraphagosomal Mycobacterium tuberculosis. Nat Med 14, 849–854 (2008).

47. Aylan B, Botella L, Gutierrez MG, Santucci P. High content quantitative imaging of Mycobacterium tuberculosis responses to acidic microenvironments within human macrophages. FEBS Open Bio 13, 1204–1217 (2023).

48. Kempker RR, et al. Lung Tissue Concentrations of Pyrazinamide among Patients with Drug-Resistant Pulmonary Tuberculosis. Antimicrob Agents Chemother 61, (2017).

49. Zhang N, et al. Optimising pyrazinamide for the treatment of tuberculosis. Eur Respir J 58, (2021).

50. Alsultan A, et al. Population Pharmacokinetics of Pyrazinamide in Patients with Tuberculosis. Antimicrob Agents Chemother 61, (2017).

51. Sambandamurthy VK, et al. A pantothenate auxotroph of Mycobacterium tuberculosis is highly attenuated and protects mice against tuberculosis. Nat Med 8, 1171–1174 (2002).

52. Sambandamurthy VK, et al. Mycobacterium tuberculosis DeltaRD1 DeltapanCD: a safe and limited replicating mutant strain that protects immunocompetent and immunocompromised mice against experimental tuberculosis. Vaccine 24, 6309–6320 (2006).

53. Lee MH, Pascopella L, Jacobs WR, Jr., Hatfull GF. Site-specific integration of mycobacteriophage L5: integration-proficient vectors for Mycobacterium smegmatis, Mycobacterium tuberculosis, and bacille Calmette-Guerin. Proc Natl Acad Sci U S A 88, 3111–3115 (1991).

54. Ehrt S, et al. Controlling gene expression in mycobacteria with anhydrotetracycline and Tet repressor. Nucleic Acids Res 33, e21 (2005).

55. Koh EI, et al. Chemical-genetic interaction mapping links carbon metabolism and cell wall structure to tuberculosis drug efficacy. Proc Natl Acad Sci U S A 119, e2201632119 (2022).

56. Pethe K, et al. A chemical genetic screen in Mycobacterium tuberculosis identifies carbon-source-dependent growth inhibitors devoid of in vivo efficacy. Nat Commun 1, 57 (2010).

57. Safi H, et al. Phase variation in Mycobacterium tuberculosis glpK produces transiently heritable drug tolerance. Proc Natl Acad Sci U S A 116, 19665–19674 (2019).

58. Gouzy A, Healy C, Schnappinger D, Ehrt SJb. Reliable detection of pyrazinamide antitubercular activity in vitro. 2022.2005. 2022.492909 (2022).

59. Hu Y, Coates AR, Mitchison DA. Sterilising action of pyrazinamide in models of dormant and rifampicin-tolerant Mycobacterium tuberculosis. Int J Tuberc Lung Dis 10, 317–322 (2006).

60. Huang Q, et al. Nutrient-starved incubation conditions enhance pyrazinamide activity against Mycobacterium tuberculosis. Chemotherapy 53, 338–343 (2007).

61. Rao M, Streur TL, Aldwell FE, Cook GM. Intracellular pH regulation by Mycobacterium smegmatis and Mycobacterium bovis BCG. Microbiology (Reading) 147, 1017–1024 (2001).

62. Berney M, et al. Essential roles of methionine and S-adenosylmethionine in the autarkic lifestyle of Mycobacterium tuberculosis. Proc Natl Acad Sci U S A 112, 10008–10013 (2015).

63. Krishnamoorthy G. Temperature jump as a new technique to study the kinetics of fast transport of protons across membranes. Biochemistry 25, 6666–6671 (1986).

64. Chen Y, Schindler M, Simon SM. A mechanism for tamoxifen-mediated inhibition of acidification. J Biol Chem 274, 18364–18373 (1999).

65. Antonenko YN, Denisov GA, Pohl P. Weak acid transport across bilayer lipid membrane in the presence of buffers. Theoretical and experimental pH profiles in the unstirred layers. Biophys J 64, 1701–1710 (1993).

66. Gutknecht J, Tosteson DC. Diffusion of weak acids across lipid bilayer membranes: effects of chemical reactions in the unstirred layers. Science 182, 1258–1261 (1973).

67. Saparov SM, Antonenko YN, Pohl P. A new model of weak acid permeation through membranes revisited: does Overton still rule? Biophys J 90, L86–88 (2006).

68. Mc DW, Tompsett R. Activation of pyrazinamide and nicotinamide in acidic environments in vitro. Am Rev Tuberc 70, 748–754 (1954).

69. Santucci P, Greenwood DJ, Fearns A, Chen K, Jiang H, Gutierrez MG. Intracellular localisation of Mycobacterium tuberculosis affects efficacy of the antibiotic pyrazinamide. Nat Commun 12, 3816 (2021).

70. Lanoix JP, Lenaerts AJ, Nuermberger EL. Heterogeneous disease progression and treatment response in a C3HeB/FeJ mouse model of tuberculosis. Dis Model Mech 8, 603–610 (2015).

71. Lanoix JP, et al. Selective Inactivity of Pyrazinamide against Tuberculosis in C3HeB/FeJ Mice Is Best Explained by Neutral pH of Caseum. Antimicrob Agents Chemother 60, 735–743 (2016).

72. Wayne LG, Hayes LG. An in vitro model for sequential study of shiftdown of Mycobacterium tuberculosis through two stages of nonreplicating persistence. Infect Immun 64, 2062–2069 (1996).

73. Kreutzfeldt KM, et al. CinA mediates multidrug tolerance in Mycobacterium tuberculosis. Nat Commun 13, 2203 (2022).

74. Gouzy A, et al. Metabolic control of drug resistance by a mycobacterial ion channel. 2026.2003.2010.710584 (2026).

75. Fontes FL, Rooker SA, Lynn-Barbe JK, Lyons MA, Crans DC, Crick DC. Pyrazinoic acid, the active form of the anti-tuberculosis drug pyrazinamide, and aromatic carboxylic acid analogs are protonophores. Front Mol Biosci 11, 1350699 (2024).

76. Bellerose MM, et al. Common Variants in the Glycerol Kinase Gene Reduce Tuberculosis Drug Efficacy. mBio 10, (2019).

77. Jain P, et al. Specialized transduction designed for precise high-throughput unmarked deletions in Mycobacterium tuberculosis. mBio 5, e01245–01214 (2014).

78. Baguet A, Koppo K, Pottier A, Derave W. Beta-alanine supplementation reduces acidosis but not oxygen uptake response during high-intensity cycling exercise. Eur J Appl Physiol 108, 495–503 (2010).

79. Vaughan RA, et al. beta-alanine suppresses malignant breast epithelial cell aggressiveness through alterations in metabolism and cellular acidity in vitro. Mol Cancer 13, 14 (2014).

80. Culviner PH, et al. Evolution of Mycobacterium tuberculosis transcription regulation is associated with increased transmission and drug resistance. Cell 188, 6620–6635 e6614 (2025).

81. Li S, et al. CRISPRi chemical genetics and comparative genomics identify genes mediating drug potency in Mycobacterium tuberculosis. Nat Microbiol 7, 766–779 (2022).

82. Kalia NP, Shi Lee B, Ab Rahman NB, Moraski GC, Miller MJ, Pethe K. Carbon metabolism modulates the efficacy of drugs targeting the cytochrome bc(1):aa(3) in Mycobacterium tuberculosis. Sci Rep 9, 8608 (2019).

83. Boshoff HI, Mizrahi V. Expression of Mycobacterium smegmatis pyrazinamidase in Mycobacterium tuberculosis confers hypersensitivity to pyrazinamide and related amides. J Bacteriol 182, 5479–5485 (2000).

84. Goude R, Roberts DM, Parish T. Electroporation of mycobacteria. Methods Mol Biol 1285, 117–130 (2015).

85. Stover CK, et al. New use of BCG for recombinant vaccines. Nature 351, 456–460 (1991).

86. Piddington DL, Kashkouli A, Buchmeier NA. Growth of Mycobacterium tuberculosis in a defined medium is very restricted by acid pH and Mg(2+) levels. Infect Immun 68, 4518–4522 (2000).

87. Gouzy A, Healy C, Black KA, Rhee KY, Ehrt S. Growth of Mycobacterium tuberculosis at acidic pH depends on lipid assimilation and is accompanied by reduced GAPDH activity. Proc Natl Acad Sci U S A 118, (2021).

88. Giske CG, Turnidge J, Canton R, Kahlmeter G, Committee ES. Update from the European Committee on Antimicrobial Susceptibility Testing (EUCAST). J Clin Microbiol 60, e0027621 (2022).

89. Johnson D, Lardy H. [15] Isolation of liver or kidney mitochondria. In: Methods in enzymology). Elsevier (1967).

90. Akerman KE, Wikstrom MK. Safranine as a probe of the mitochondrial membrane potential. FEBS Lett 68, 191–197 (1976).

91. Mueller P, Rudin DO, Tien HT, Wescott WCJTjopc. Methods for the formation of single bimolecular lipid membranes in aqueous solution. 67, 534–535 (1963).

92. Denisov SS, Kotova EA, Khailova LS, Korshunova GA, Antonenko YN. Tuning the hydrophobicity overcomes unfavorable deprotonation making octylamino-substituted 7-nitrobenz-2-oxa-1,3-diazole (n-octylamino-NBD) a protonophore and uncoupler of oxidative phosphorylation in mitochondria. Bioelectrochemistry 98, 30–38 (2014).

